# Molecular and morphological analyses support recognition of *Prostanthera volucris* (Lamiaceae), a new species from the Central Tablelands of New South Wales

**DOI:** 10.1101/2021.12.21.473648

**Authors:** Ryan P. O’Donnell, Jeremy J. Bruhl, Ian R.H. Telford, Trevor C. Wilson, Heidi C. Zimmer, Guy M. Taseski, Rose L. Andrew

## Abstract

Research into the systematics of *Prostanthera* has recently revealed a close evolutionary relationship among *P. phylicifolia s. str.*, the critically endangered *P. gilesii*, and a population of uncertain identity from the Central Tablelands of New South Wales, Australia. Previous analyses were unable to establish whether genetic boundaries separated these taxa. This study aimed to assess the species boundaries among these three taxa using a combination of single-nucleotide polymorphisms (SNP) sampled at the population-scale and multivariate analysis of morphological characters. Non-parametric and parametric statistics, neighbour-network analysis, phylogenetic analysis, and ancestry coefficient estimates all provided support for discrete genetic differences between the three taxa. Morphological phenetic analysis identified a suite of characters that distinguished each of these taxa. This corroboration of evidence supports the presence of three independently evolving lineages. *Prostanthera gilesii* and *P. phylicifolia s. str.* are distinct species independent from the third taxon which is described here as *P. volucris* R.P.O’Donnell. A detailed description, diagnostic line drawings and photographs are provided. We evaluate *P. volucris* as satisfying criteria to be considered Critically Endangered.

## Introduction

*Prostanthera* Labill. is the most speciose genus of endemic Australian Lamiaceae, encompassing over 105 accepted species (Conn 1984, 1988; APC 2021; Conn et al. 2021). Recent studies have revealed the genus to be far more diverse than currently recognised, with several species complexes that require resolution (Wilson et al. 2012; Conn et al. 2016; Wilson et al. 2019; Conn et al. 2021; O’Donnell et al. 2021). Many species of *Prostanthera* are niche specialists and tend to grow on isolated, rocky outcrops (Conn 1984, 1988; Conn & Wilson 2015b; Conn & Wilson 2015a; Wilson et al. 2019). Like other rocky outcrop specialists, these species have highly restricted ranges and are particularly vulnerable to environmental threats (Fitzsimons & Michael 2017; Selwood & Zimmer 2020; Hopper et al. 2021; Silveira et al. 2021). Among its accepted species, 19 are listed as threatened at a Commonwealth level, representing almost a fifth of the genus (Department of Agriculture 1999a, 2021). One example is the critically endangered *Prostanthera gilesii* G.W.Althofer *ex* B.J.Conn & T.C.Wilson, which is geographically restricted in comparison with its close relative, *P. phylicifolia* F.Muell.

*Prostanthera gilesii* is currently the subject of a targeted conservation and management project (OEH 2019; Scott & Auld 2020; Andrew et al. unpublished data). The species is currently known only from two small subpopulations within the Mount Canobolas State Conservation Area, south-west of Orange in the Central Tablelands of New South Wales (Conn & Wilson 2015b). Molecular phylogenies recovered a close relationship between *P. gilesii* and *P. phylicifolia s. str*., the latter composed of populations distributed across the Victorian Alps and Snowy Mountains, Monaro, and Southern Tablelands of New South Wales (O’Donnell et al. 2021). Another population occurring at the Evans Crown Nature Reserve (hereafter referred to as *P.* sp. Evans Crown), located approximately 2.8 km south-east of Tarana, New South Wales, was recovered as sister to *P. gilesii.*

While superficially similar to *P. phylicifolia* and *P. gilesii*, *P.* sp. Evans Crown is geographically disjunct and differs substantially with respect to several morphological characters (O’Donnell et al. 2021). These taxa also differ in the substrates they occupy; *P. phylicifolia* and *P.* sp. Evans Crown are restricted to granite outcrops, while *P. gilesii* is found on basaltic substrates (NPWS 2009; OEH 2019; Scott & Auld 2020). These differences are not encompassed by diagnoses of *P. phylicifolia* or *P. gilesii* (Mueller 1858; Conn & Wilson 2015b).

As phylogenetic data produced variable topologies with respect to these three taxa, there was insufficient evidence to determine whether genetic boundaries separate them (O’Donnell et al. 2021). The aim of this study was to elucidate the relationship between *P. gilesii*, *P. phylicifolia,* and *P.* sp. Evans Crown, and to assess whether they each represent independently evolving lineages that warrant species level recognition. Each taxon was sampled at the population scale to produce reduced-representation single nucleotide polymorphism (SNP) data, which were then analysed using population genetic and phylogenetic methods. Morphometric phenetic analyses were then applied to objectively assess interspecific morphological variation and distinguish informative characters for identification.

## Materials and Methods

### Integrative taxonomic approach

To recover a stable taxonomic classification that agrees with evolutionary history, and to avoid the possibility of taxonomic over-inflation, an integrative taxonomic approach must be applied to questions of species delimitation in *Prostanthera*. Integrative taxonomy aims to incorporate and synthesise multiple, independent lines of evidence to rigorously test and corroborate delimitation hypotheses, thereby increasing the probability that a set of independently evolving metapopulation lineages—i.e., species, *sensu* de Queiroz (2007)— will be resolved (Dayrat 2005; Schlick-Steiner et al. 2009; Padial et al. 2010). Schlick-Steiner et al. (2009) provide a framework for an integrative taxonomic approach, suggesting that morphology be used as a primary line of evidence, followed by a genetic discipline. Previous studies of *Prostanthera* have incorporated phenetic analyses of morphological characters to rigorously assess phenotypic variation between putative taxa (Conn 1984; Conn et al. 2013; Wilson et al. 2017; Conn et al. 2021), and the use of these techniques is widely accepted. Previous molecular studies of *Prostanthera* incorporating Sanger sequencing data have provided some species-level resolution (Wilson et al. 2012; Conn et al. 2013; Conn et al. 2016; Conn et al. 2021; O’Donnell et al. 2021). Phylogenetic analysis by O’Donnell et al. (2021), however, recovered discordant topologies between nuclear and chloroplast datasets, suggesting that hybridisation, introgression, or incomplete lineage sorting may have occurred within *Prostanthera*. The use of high-throughput sequencing molecular approaches capable of detecting genome-wide admixture was recommended for future studies of *Prostanthera* to mitigate the confounding effect of these demographic processes (O’Donnell et al. 2021).

### Genotyping-by-sequencing

DArTseq (Kilian et al. 2012), is a cost-competitive reduced-representation sequencing (RRS) platform that has provided resolution and demographic inference at the population scale in taxonomically recalcitrant plant groups (Sansaloni et al. 2010; Steane et al. 2011). Reduced-representation sequencing allows for a targeted fraction of an organism’s genome to be sequenced in species with little pre-existing reference information (Narum et al. 2013; Soltis et al. 2013; Fernández-Mazuecos et al. 2017). In contrast to microsatellite or amplified fragment length polymorphism (AFLP) approaches which rely only on a small number of neutral molecular markers representing a limited subset of the genome, RRS approaches greatly increase the number of neutral markers assayed, thereby improving the precision of estimates of population structure, admixture, and demography (Narum et al. 2013). DArTseq captures SNP data and recent studies at the population scale have been able to identify distinct genetic lineages in morphologically variable groups that warrant specific recognition (Joyce et al. 2021). DNA extraction, library preparation and genotyping-by-sequencing using the DArTseq platform was conducted by Diversity Arrays Technology (DArT) P/L (Canberra, Australia). The resulting reads were filtered, assembled *de novo* and scored by DArT internally using their proprietary analysis pipeline following methods similar to those outlined by Kilian et al. (2012) and Cruz et al. (2013).

### Sampling

All populations of *P. gilesii, P. phylicifolia* and *P.* sp. Evans Crown sampled by O’Donnell et al. (2021) were used again in this study to closely replicate their sampling strategy. Material was sourced from herbarium specimens or field collections of silica-dried or freeze-dried leaf material. *Prostanthera phylicifolia* was collected from 12 populations spanning its geographic distribution, *P. gilesii* was collected from both known subpopulations within the Mount Canobolas State Conservation Area (here referred to as Towac and Walls), and *P.* sp. Evans Crown was collected from within the Evans Crown Nature Reserve (Figure 1). Since the compact and tangled habit of *P. gilesii* and *P.* sp. Evans Crown complicated an accurate estimation of individual composition, samples were collected from individuals at least 6 m apart to reduce repeated sampling of the same genet where possible (e.g. the Walls population appears to be two plants within a shared 3 m diameter space). Herbarium vouchers were lodged at either or both the National Herbarium of New South Wales (NSW) or the N.C.W. Beadle Herbarium (NE). Flowering material was collected and preserved in 70% ethanol for morphological examination.

**Fig. 1.**
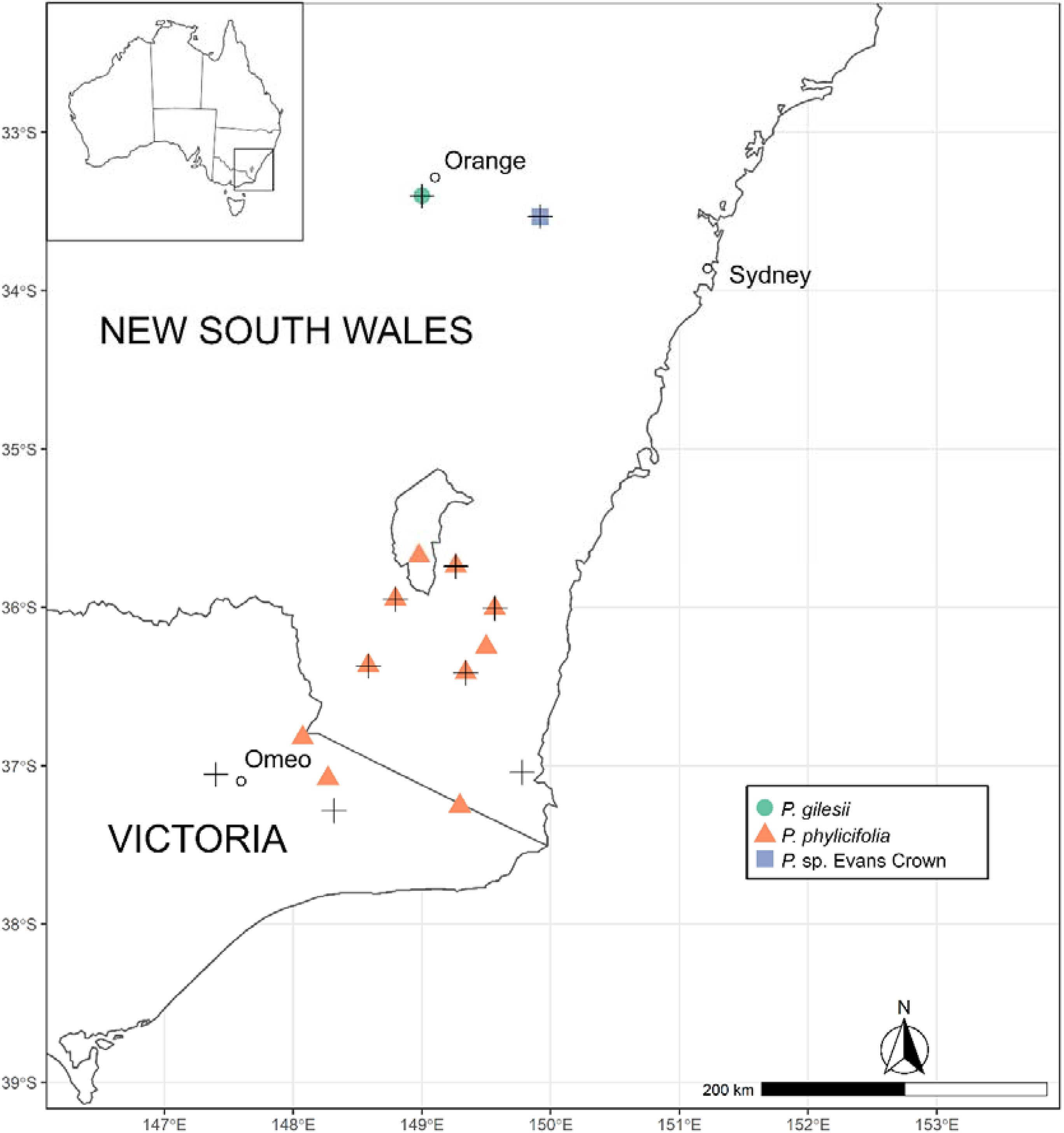
Distribution of *Prostanthera gilesii*, *P. phylicifolia* and *P.* sp. Evans Crown populations sampled in this study. Populations with associated herbarium vouchers that were measured for morphological phenetic analysis (Online Resource 2) are identified with shapes, while crosses indicate populations that were sampled for genomic analysis (Online Resource 1)

To examine population dynamics and genetic boundaries between closely related taxa, 38 samples comprising 30 samples of *P.* sp. Evans Crown, seven samples of *P. phylicifolia* and one sample of *P. gilesii* (Walls) were newly sequenced for this study to produce SNP data. New samples were co-analysed with samples of *P. gilesii* (65)*, P. phylicifolia* (13), and *P.* sp. Evans Crown (7) that had been previously sequenced as part of targeted conservation and management projects underway for *P. gilesii* (OEH 2019; Andrew et al. unpublished data). One sample of *P. phylicifolia* that had previously been sampled as part of a targeted conservation and management project for *P. densa* A.A. Ham. and *P. marifolia* R.Br. was included in this co-analysis (ReCER 2020). To ensure datasets were compatible for co-analysis, three samples from the preliminary *P. gilesii* SNP dataset were sequenced again as technical duplicates (Online Resource 1). In total, 125 samples were included in this co-analysis (Online Resource 1).

To provide outgroup representatives for phylogenetic analysis, an additional SNP dataset was produced where selected samples of *P. gilesii* (4)*, P. phylicifolia* (14) and *P.* sp. Evans Crown (4) were co-analysed with samples of *P. densa* (2)*, P. marifolia* (3), *P. granitica* Maiden & Betche (2) and *P. scutellarioides* (R.Br.) Briq. (2) that had previously been sampled as part of a targeted conservation and management project for *P. densa* and *P. marifolia* (ReCER 2020). These species were chosen as outgroup representatives as they were previously recovered as sister clades to the clade containing *P. gilesii*, *P. phylicifolia*, and *P.* sp. Evans Crown (O’Donnell et al. 2021). In total, 31 samples were included for this analysis.

For morphometric phenetic analysis, 20 herbarium voucher specimens were measured and scored (Figure 1, Online Resource 2). For both *P. gilesii* and *P.* sp. Evans Crown, five voucher specimens were scored. For *P. phylicifolia*, 10 specimens were scored to cover the species’ broader distribution. Each herbarium voucher specimen was considered as an individual plant and OTU when scoring. Voucher specimens were selected to incorporate the extent of variability within a taxon, and on the basis that they could provide three replicates for each character being scored.

### Molecular analysis of SNP data

The SNP dataset delivered by DArT was processed with the package *dartR* (Gruber et al. 2018) in the statistical package R (Team 2013). Processing followed methods outlined by Gruber et al. (2018). A random member of each pair of technical duplicates was first removed, and sites were then filtered to require 99% repeatability (as estimated by the DArT proprietary pipeline). Monomorphic loci were then removed, and the remaining loci were filtered based on a minimum call rate of 95%. Loci representing secondary SNPs (i.e. sequenced fragments with more than one SNP) were then removed. Following site filtering, individuals were then filtered based on individual call rate, requiring a minimum call rate of 90%. Loci with similar trimmed sequence tags were then removed (threshold = 25%). The resultant dataset comprised 9,691 loci scored for 114 individuals, i.e. 11 individuals were removed by filtering (Online Resource 1). Filtering methods were the same for the secondary dataset used for phylogenetic analysis, with the exception of filters for loci and individual call rates. The minimum locus call rate was lowered to 80% to keep as many loci as possible and the minimum individual call rate was lowered to 60% to ensure outgroup representatives were not filtered out. Three individuals were removed by filtering and the resultant dataset for phylogenetic analysis comprised 2,095 loci scored for 28 individuals (Online Resource 1).

To assess the similarities between populations and individuals, ordination of the dataset using principal coordinates analysis (PCoA) was conducted using the ‘gl.pcoa’ function in *dartR* (Gruber et al. 2018). To perform a neighbour-network analysis and visualise a distance-based network, a Euclidian distance matrix was calculated using the ‘gl.dist.ind’ function in *dartR* (Gruber et al. 2018) and exported to SplitsTree5 (Huson & Bryant 2006) using the ‘splitstree’ function in *RSplitsTree* (Bickel & Zakharko 2016).

To examine phylogenetic relationship, trees were estimated from the concatenated SNP matrix under a coalescent model using SVDquartets (Chifman & Kubatko 2014, 2015) as implemented in PAUP* v4.0a169 (Swofford 2002). Quartets were sampled exhaustively using the Quartet FM (QFM) quartet assembly algorithm, with 1,000 multilocus bootstrap replications conducted to estimate branch support. Bootstrap support values were considered strong if they provided support values ≥95%, moderate from 80–94% and weak from 50– 79%. The final best-scoring tree was visualised using TreeGraph 2 (Stöver & Müller 2010).

Exclusion of clones and close relatives helps avoid bias in population genetic parameters in species with mixed sexual and asexual reproduction. To assess genomic relationships between sampled individuals, genomic relationship matrices were calculated for each putative taxon using the ‘stamppGmatrix’ function in the R package *StAMPP* (Pembleton et al. 2013) which calculates genomic relationships following methods described by Yang et al (2010). To mitigate the potential confounding effect of the presence of clonal individuals in calculations of population statistics, subsequent analyses were run using a subset of samples comprised of individuals thought to confidently represent genets (Online Resource 1). Individuals were selected by visual inspection of the genomic relationship matrices and associated dendrograms. An individual from each distinct cluster in the dendrogram was chosen. Clusters of samples that visually resembled unresolved polytomies in the dendrogram were considered to represent possible clones and only one sample per polytomy was included in further analyses. As most samples of *P. gilesii* (Towac) appeared to represent possible clones, field observations were used to identify individuals that appeared most likely to represent distinct genets on the basis of physical distance and separation. Estimates of heterozygosity and *F*_*ST*_ involving that population should therefore be interpreted with caution. Inspection of genomic relationship matrices resulted in the exclusion of 58 samples of *P. gilesii* and 28 samples of *P.* sp. Evans Crown.

To assess levels of genetic diversity, observed heterozygosity (*H*_*O*_) and expected heterozygosity (*H*_*E*_) values were calculated using the ‘gl.report.heterozygosity’ function in *dartR.* Statistics were calculated at putative species level based on clusters observed in PCoA and neighbour-network analyses. Statistics were also calculated for populations with five or more samples. To assess levels of genetic divergence, pairwise *F*_*ST*_ values were calculated for putative species groups, and between populations with five or more samples using the ‘stamppFst’ function in the R package *StAMPP* (Pembleton et al. 2013) which estimates pairwise *F*_*ST*_ values according to Weir and Cockerham (1984). To estimate statistical support, 1000 bootstrap replicates were conducted based on a 95% confidence interval. Both subpopulations of *P. gilesii* were treated as a single population in these calculations. Other populations with five or more samples that were used in calculations included the Dangelong Nature Reserve and Adaminaby populations of *P. phylicifolia*, and the Evans Crown population. *F*_*ST*_ values <0.05 were considered to indicate low genetic differentiation, 0.05– 0.25 indicated moderate genetic differentiation, and values >0.25 indicated pronounced differentiation (Freeland et al. 2011).

To investigate admixture, individual ancestry coefficients were estimated using Sparse Non-Negative Matrix Factorisation (sNMF) in the R package *LEA* (Frichot & François 2015). The package uses cross-entropy values to infer the probable number of ancestral populations (*K*) in the data and assign individuals to genetic clusters and the optimal number of ancestral populations was selected based on the post-stabilisation of the steepest decline in cross-entropy values (Frichot & François 2015; van der Merwe et al. 2021). The ‘snmf’ function was executed for values of *K* = 1–8 with 50 replicates for each value of *K*.

### Phenetic analysis of morphological data

The character list for morphometric phenetic analysis (Online Resource 3) consisted of 29 morphological characters, comprising 12 vegetative and 17 reproductive characters. Characters were selected from previous morphological studies of *Prostanthera* (Conn 1984; Williams et al. 2006; Conn et al. 2013), with additional characters added following preliminary examination of specimens (i.e. indumentum direction and density). The mean of replicates (three per specimen) for quantitative characters (23 in total) was used in morphometric analysis.

Reproductive characters only included calyx, prophyll and podium characters and did not include androecial or gynoecial characters as an insufficient number of specimens exhibited useful reproductive material to score such characters consistently. The majority of measurements were made from herbarium vouchers, except for three specimens of *P.* sp. Evans Crown (Taseski 853, O’Donnell 30, O’Donnell 55) and one specimen of *P. phylicifolia* (O’Donnell 61), where calyx and prophyll characters were scored from flowers preserved in 70% ethanol. The final matrix contained 29 characters scored for 20 specimens, including 23 quantitative and 6 qualitative characters.

The morphological data matrix was analysed in PATN 4.0 (Belbin & Collins 2013) using default settings. All characters were weighted equally in each analysis with the Gower association metric, as it is suited to datasets that contain a combination of quantitative, binary, and qualitative characters (Sokal 1986). The unweighted pair-group method using arithmetic means (UPGMA) was used as a means for cluster analysis as it is considered to accurately represent real distances between individuals in taxonomic datasets (Chappill & Ladiges 1992; Copeland et al. 2007). Semi-strong hybrid multidimensional scaling (SSH-MDS) was used for ordination analyses as it has been shown to accurately reflect phenetic patterns (Minchin 1987) and has frequently been used in recent morphological phenetic studies, including studies of *Prostanthera* (Plunkett et al. 2009; Conn et al. 2013; Bean 2014; de Salas & Schmidt-Lebuhn 2018).

## Results

### Principal component analysis of SNP data

PCoA ordination of SNP data (Figures 2a–b) organised samples as three general clusters: 1) all samples of *P.* sp. Evans Crown as a single cluster; 2) all samples of *P. gilesii*, subdivided into two subclusters corresponding to their respective subpopulation of origin; and 3) all sampled populations of *P. phylicifolia*, subdivided into subclusters corresponding to their respective population of origin.

**Fig. 2.**
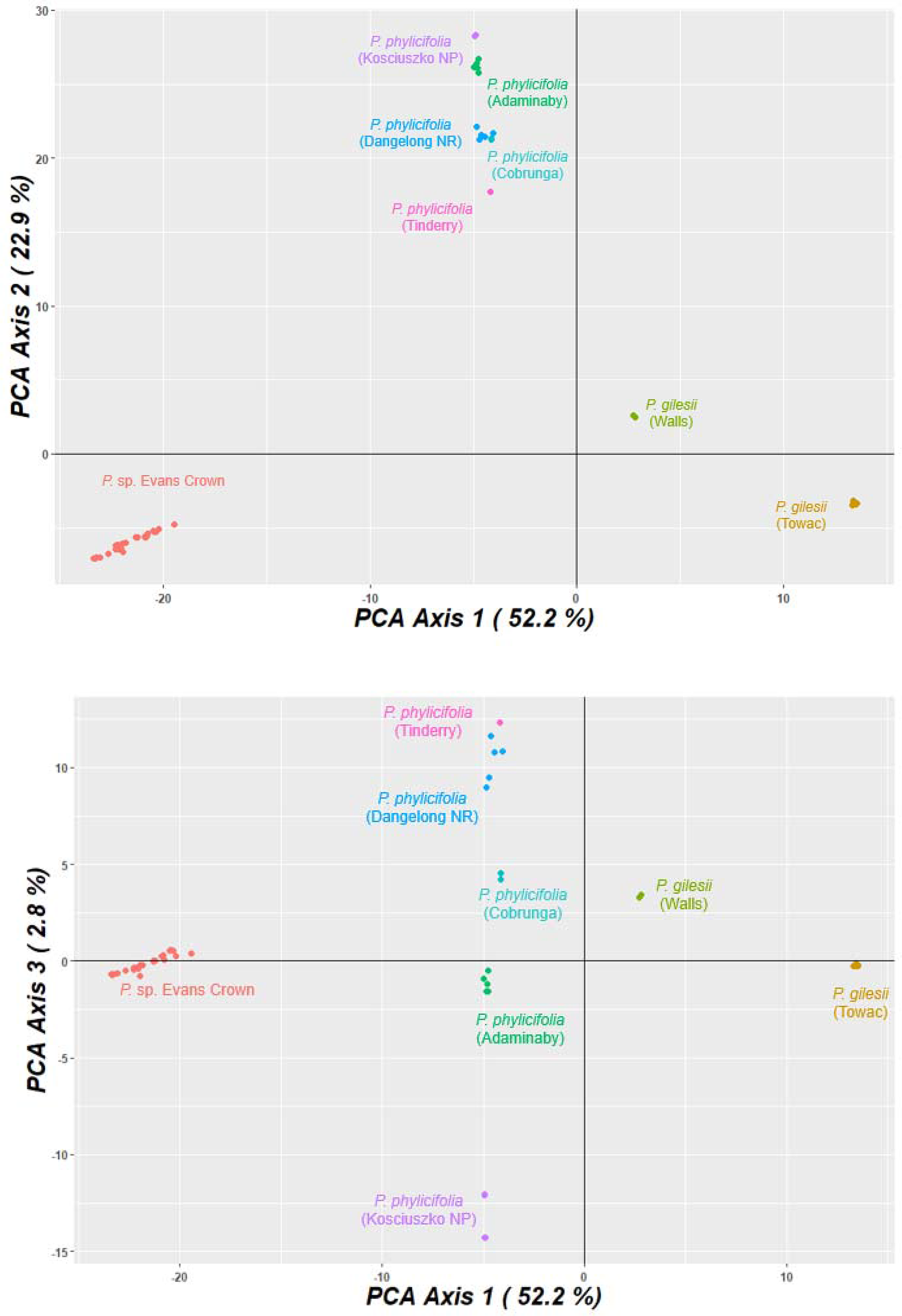
**a–b** Principal coordinate analysis (PCoA) plot of DArTseq SNP data from all sampled individuals and populations. 2a = PC1vPC2; 2b = PC1vPC3. NP = National Park, NR = Nature Reserve

### Neighbour-network analysis

The network graph (Figure 3) showed three main branches exhibiting little to no reticulation between them. One branch included all samples of *P.* sp. Evans Crown with some reticulation present among individuals. Another branch included all samples of *P. phylicifolia* which exhibited limited reticulation among individuals, but some reticulation between populations. Reticulation was observed within two main groups: one consisting of individuals from Tinderry and Dangelong Nature Reserve; and another consisting of individuals from Adaminaby, Kosciuszko National Park and Cobrunga. The third branch included all individuals of *P. gilesii*, with both specimens from the Walls population forming a distinct subbranch. Little to no reticulation was present among individuals or between the two sub-branches of *P. gilesii*.

**Fig. 3.**
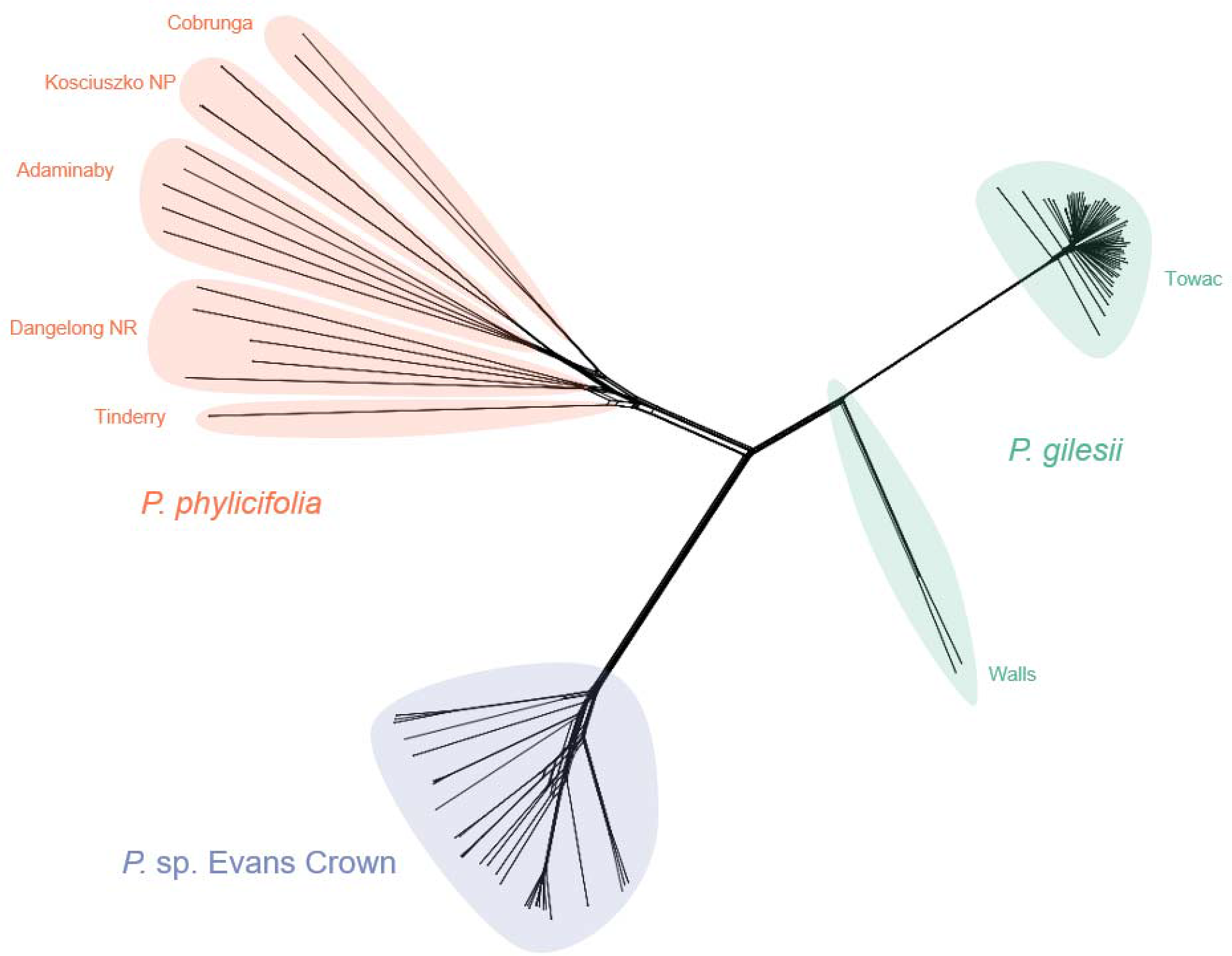
Neighbour-network graph produced by SplitsTree5 of DArTseq SNP data from all sampled individuals and populations. Putative species groups are coloured, and populations are labelled. NP = National Park, NR = Nature Reserve

### Phylogenetic analysis

Exhaustive quartet sampling sampled 20,475 quartets in total, with a compatible quartet weight of 89.47% (18,318 quartets). The final assembled tree (Figure 4) was comprised of two core clades. One strongly supported clade (BS = 100%) contained all ingroup samples of *P. gilesii, P. phylicifolia* and *P.* sp. Evans Crown, while the other contained all outgroup representatives. Within the ingroup clade, all samples of *P. phylicifolia* were recovered as a moderately supported clade (BS = 85%), and all samples of *P. gilesii* and *P.* sp. Evans Crown were recovered within a strongly supported clade (BS = 100%). Within this clade, all samples of *P. gilesii* formed a strongly supported clade (BS = 100%) and all samples of *P.* sp. Evans Crown formed a strongly supported clade (BS = 100%).

**Fig. 4.**
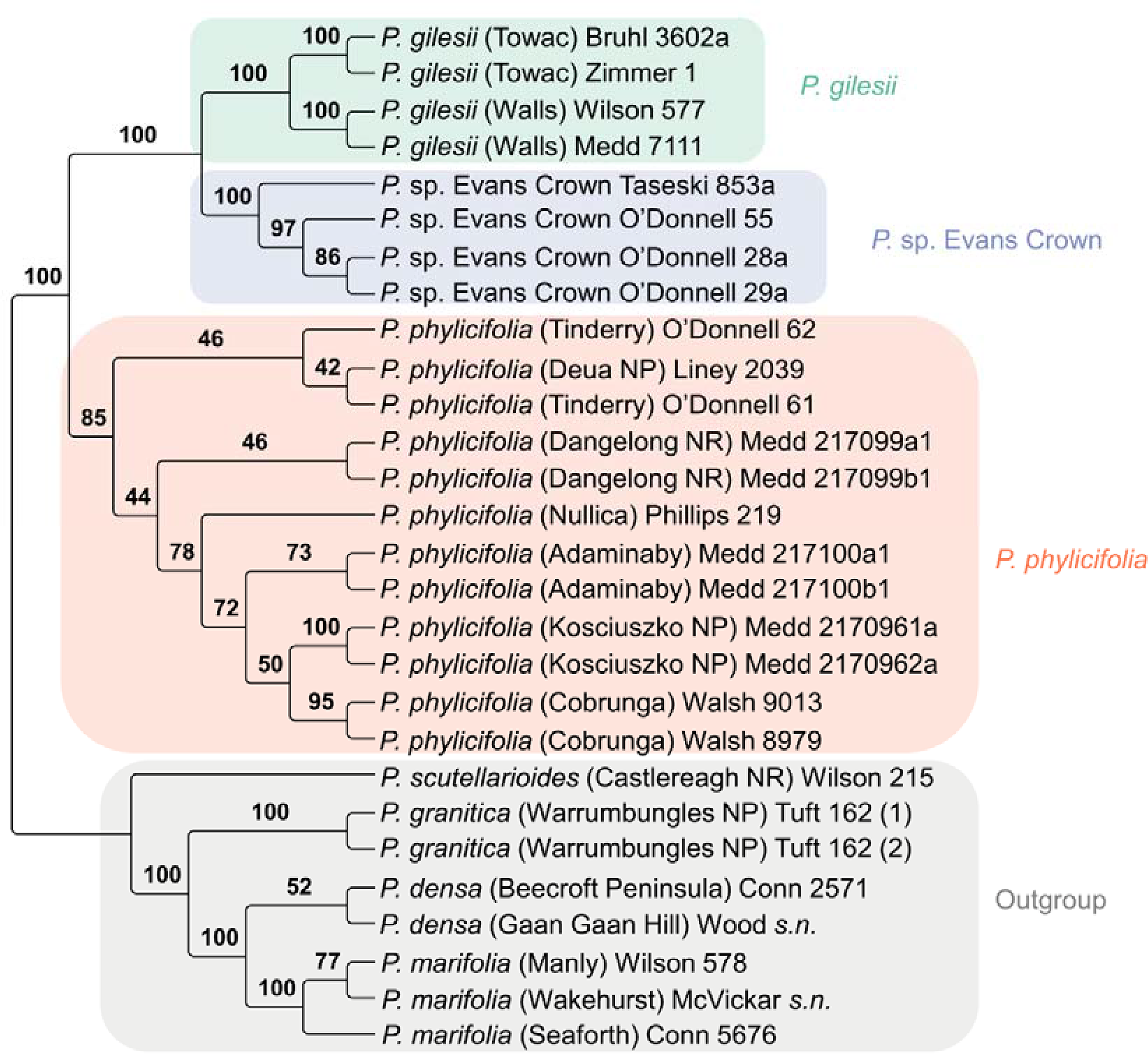
Phylogeny generated by SVDQuartets analysis of DArTseq SNP data for 28 samples of *Prostanthera*. Putative species groups are coloured, and populations are labelled. Labels are species/phrase names and population of origin, followed by primary collector and collection number

### sNMF cluster analysis

Based on flattening of the cross-entropy curve (Online Resource 4), *K* = 3 was the most likely number of ancestral populations represented in the data. Estimates of ancestry coefficients for *K =* 3 recovered all samples of *P. phylicifolia* as one cluster, all samples of *P.* sp. Evans Crown as a second cluster, and all samples of *P. gilesii* as a third cluster (Figure 5). Each cluster exhibited little to no shared ancestry with other clusters. Because of the small number of individuals in the Walls subpopulation of *P. gilesii*, it is not possible to evaluate whether this population may or may not represent a distinct genetic cluster. These results therefore must be interpreted with caution. To compare alternative delimitation models, results for *K =* 3 were compared with results for *K =* 2 and *K =* 4. For *K =* 2, all samples of *P. gilesii* formed a cluster, and all samples of *P.* sp. Evans Crown formed a cluster. Samples of *P. phylicifolia* were recovered as individuals with shared ancestry from both clusters. For *K* = 4, samples of *P. phylicifolia* were grouped into two clusters; the first comprised of samples from Tinderry, Dangelong Nature Reserve, Cobrunga and Adaminaby, and the second comprised of samples from Kosciuszko National Park.

**Fig. 5.**
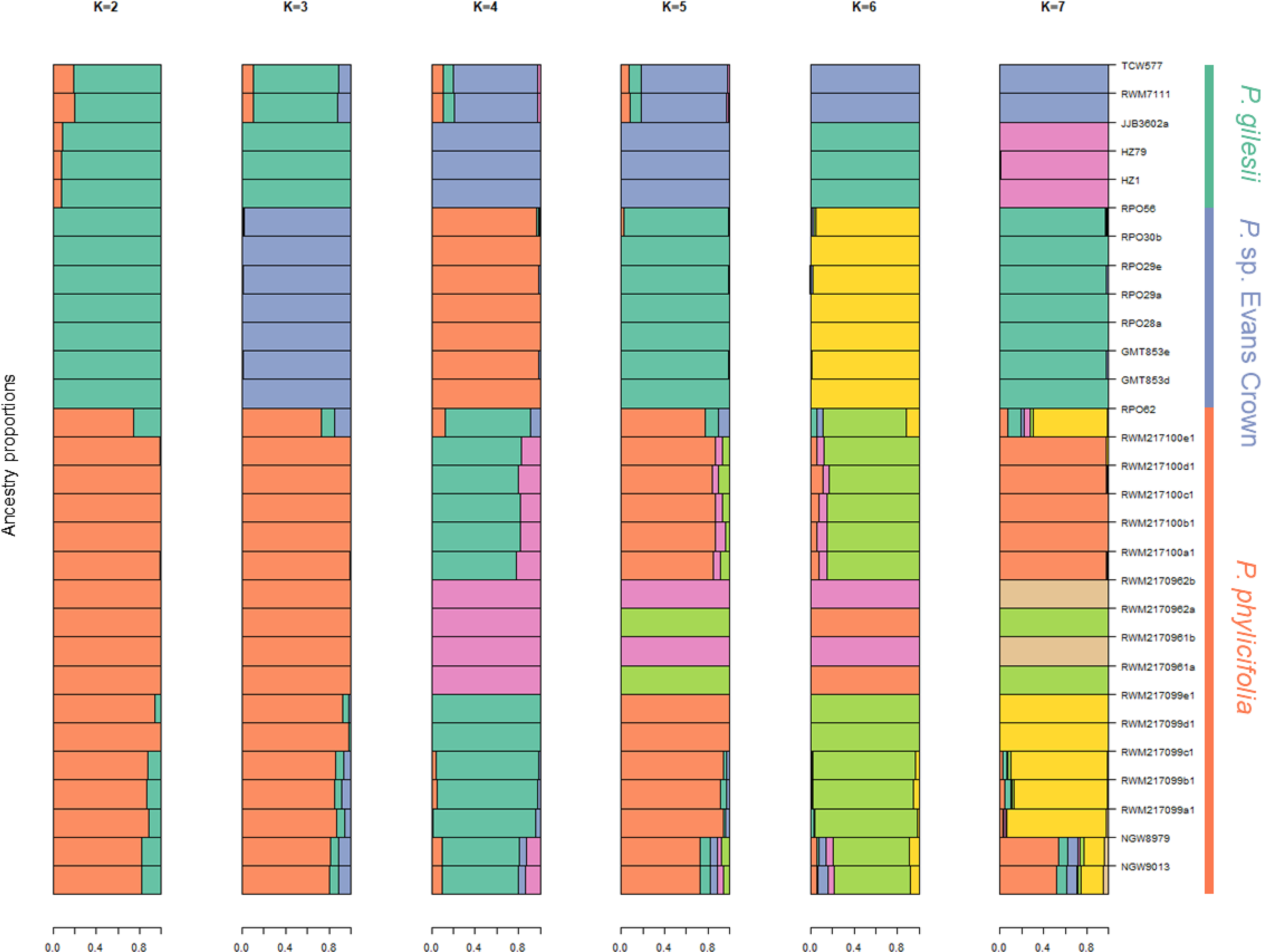
Individual ancestry proportions from model-based clustering using sNMF of all sampled individuals for values of K = 2–7. Putative species groups are labelled, and black lines have been added to demarcate individuals. Sample codes follow those outlined in Online Resource 1

### Population genetic analysis

Genomic relationship matrices (Online Resources 5–7) indicated little differentiation between specimens of *P. gilesii*, while *P.* sp. Evans Crown and *P. phylicifolia* exhibited more diversity among individuals. Following removal of putative clones, the highest population-scale *F*_*ST*_ value of 0.638 was observed between *P.* sp. Evans Crown and *P. gilesii,* and between *P.* sp. Evans Crown and *P. phylicifolia,* while the lowest *F*_*ST*_ value (*F*_*ST*_ = 0.166) was between the two populations of *P. phylicifolia* (Online Resource 8). Observed and expected heterozygosity were highest in the Dangelong population of *P. phylicifolia* (*H*_*O*_ = 0.170; *H*_*E*_ = 0.170) and lowest in *P.* sp. Evans Crown (*H*_*O*_ = 0.043; *H*_*E*_ = 0.043) (Table 1). Species-level results (Table 2; Online Resource 9) were concordant with those at the population level.

**Table 1.**
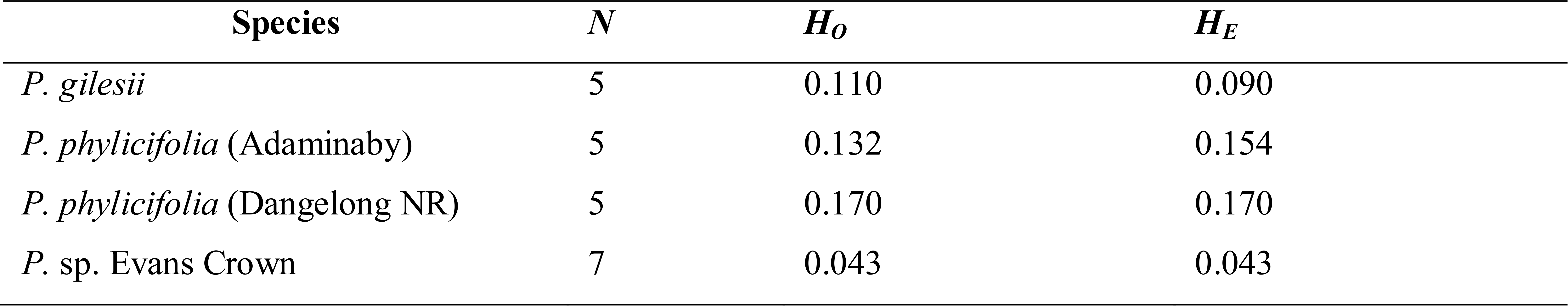
Basic summary statistics for populations of *P. gilesii, P. phylicifolia* and *P.* sp. Evans Crown where *N* ≥ 5, following removal of putative clones

**Table 2.**
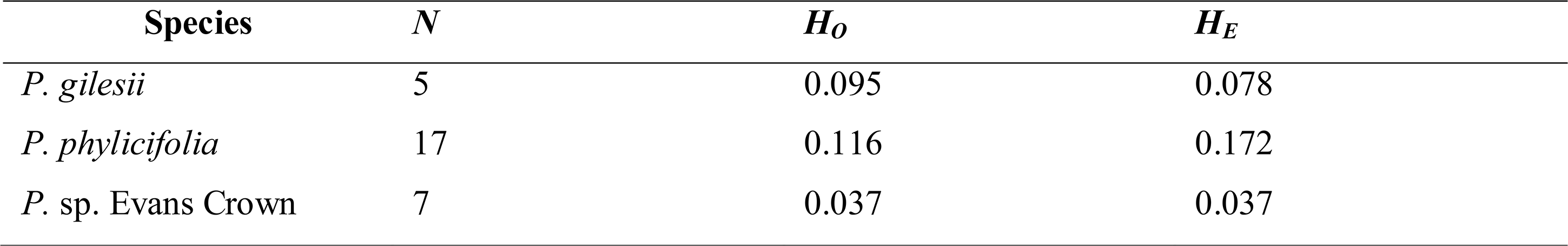
Basic summary statistics for *P. gilesii, P. phylicifolia* and *P.* sp. Evans Crown following removal of putative clones

### Morphometric analysis

Cluster analysis recovered three groups at a dissimilarity value of 0.2233 (Figure 6a). The first group consisted of all specimens of *P.* sp. Evans Crown, the second consisted of all specimens of *P. gilesii*, and the third consisted of all specimens of *P. phylicifolia.* The top five characters that contributed to the separation of these groups based on Kruskal-Wallis values were lamina width, branch hair density, petiole hair direction, prophyll hair density and lamina length (Table 3).

**Fig. 6.**
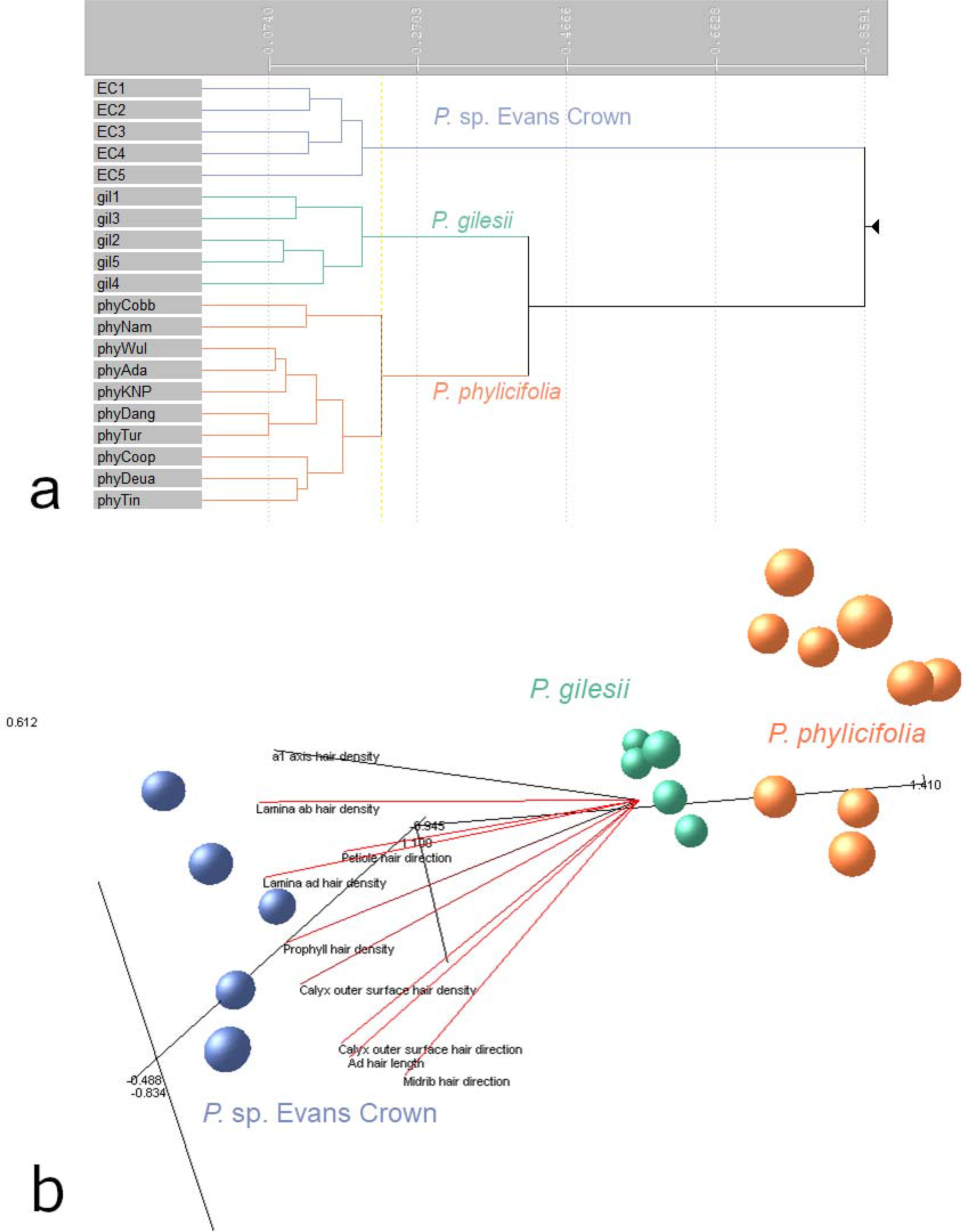
**a–b** Output from morphometric analysis of 29 characters measured from 20 specimens of *Prostanthera* showing three discrete groups. a = flexible unweighted pair-group method with arithmetic mean (UPGMA) phenogram; b = semi-strong hybrid multidimensional scaling (SSH-MDS) ordination with characters with PCC vectors with *r*^*2*^ values >0.9 (stress = 0.0503). See Online Resource 2 for OTU codes, Online Resource 3 for character list, and Online Resource 10 for PCC values

**Table 3.**
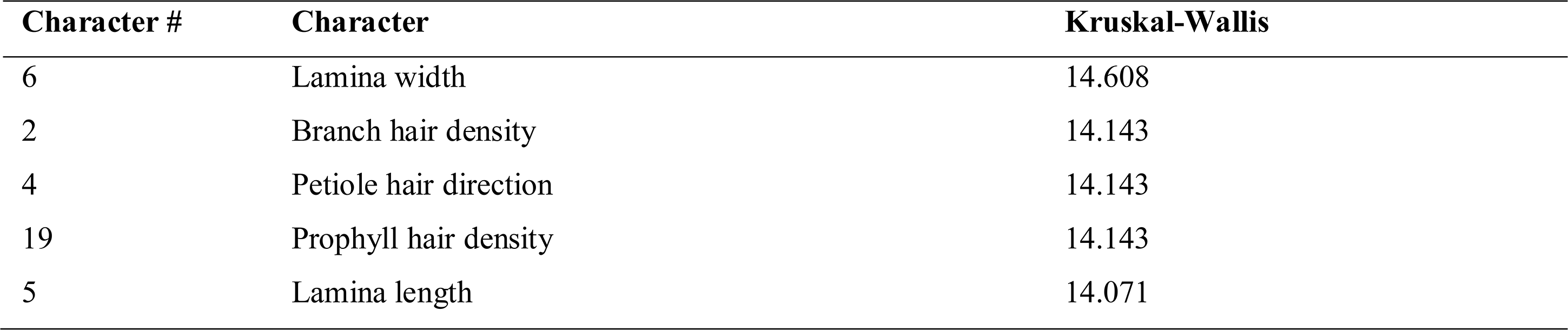
Top five characters contributing to distinction between groups in the phenetic cluster analysis in Fig. 4 based on Kruskal-Wallis (KW) values

Three discrete clusters were observed in ordination plots: one cluster contained all specimens of *P.* sp. Evans Crown; another contained all samples of *P. gilesii* and the remaining cluster contained all samples of *P. phylicifolia* (Figure 6b). The stress value of 0.0503 is interpreted as low, suggesting that the ordination is an accurate representation of the dataset in reduced dimensionality (Belbin 2013). Petiole hair direction and prophyll hair density, which were shown to be informative characters based on Kruskal-Wallis values in the cluster analysis, were also recovered with *r*^*2*^ values >0.9 for the ordination (Online Resource 10).

## Discussion

Corroborating molecular (Figures 2–5; Online Resources 8, 9) and morphological (Figure 6) lines of evidence support close relationships among *P. gilesii, P. phylicifolia* and *P.* sp. Evans Crown established by O’Donnell et al. (2021) and demonstrate that they each represent independently evolving lineages.

Phenetic analyses of SNP data (Figure 2) recovered three core clusters congruent with the two named and one putative species. Both the neighbour-network tree (Figure 3) and sNMF ancestry coefficient estimate plots (Figure 5) demonstrated little reticulation or recombination among the three groups, indicating that gene flow between these groups is highly restricted. While phylogenies recovered in this study (Figure 4) and by O’Donnell et al. (2021) both indicate a sister relationship between *P. gilesii* and *P.* sp. Evans Crown, pairwise *F*_*ST*_ values between the two (Online Resource 9) suggest that they are more genetically different from one another than either is to *P. phylicifolia*, despite the relatively small geographic distance between *P. gilesii* and *P.* sp. Evans Crown. The observed heterozygosity values reported (Tables 1, 2) suggest that accelerated genetic drift in historically small populations has likely contributed to the high *F*_*ST*_ values reported for *P. gilesii* and *P.* sp. Evans Crown. Nevertheless, the values reported here indicate substantial genetic divergence at the population and putative species level.

The phylogenies presented by O’Donnell et al. (2021) recovered two distinct clades of *P. phylicifolia s. str.*; a ‘western’ clade comprised of populations occurring along the Victorian Alps and Snowy Mountains, and an ‘eastern’ clade comprised of populations from Tinderry south to Dangelong. The phylogeny estimated in this study (Figure 4) placed all samples of *P. phylicifolia* within a monophyletic clade, but did not recover the same eastern and western clades. The neighbour-network analysis (Figure 3) recovered branches that had similar membership to the eastern and western clades recovered by O’Donnell et al. (2021), but both branches exhibited moderate amounts of reticulation, suggesting that gene flow has occurred between the two groups. This may account for the relatively low support values within the *P. phylicifolia* clade seen in the phylogenetic analysis in this study (Figure 4). PCoA ordination of SNP data (Figure 2) recovered several subclusters that corresponded with the sample’s population of origin, within a larger cluster containing all specimens of *P. phylicifolia.* Morphological analyses in this study (Figure 6) did not detect substantial variation between populations of *P. phylicifolia*; however, clusters recovered within this group reflected similar patterns to results recovered by O’Donnell et al. (2021). For example, specimens from Dangelong Nature Reserve, Tuross Falls, Mt. Coopracambra, Tinderry and Deua National Park formed two clusters corresponding with the eastern clade while samples from Wulgulmerang, Adaminaby, and Kosciuszko National Park clustered together, corresponding with the western clade.

While two separate clades of *P. phylicifolia s. str.* were recovered by O’Donnell et al. (2021), the presence of gene flow and lack of substantial morphological differentiation demonstrated here indicate that there is likely no major distinction between these clades. It is possible that these clades represent lineages that are either in the process of diverging, or conversely, lineages that were once geographically isolated that have become reconnected. From a conservation perspective it is important to recognise that these clades are present, as they may represent lineages adapted to variable habitats. Preservation of these lineages may aid the resilience of this species in light of anticipated future climate change. Additional population-scale sampling, particularly of populations that appear to be intermediary between the eastern and western clades (e.g. Nullica State Forest), will further our understanding of the structure of phenotypic and genetic variation within this species. All populations share a cohesive morphology, which is congruent with Mueller’s (1858) protologue and type localities. Our results therefore support the recognition of populations from the Victorian Alps and Snowy Mountains, Monaro, and Southern Tablelands of New South Wales as *P. phylicifolia s. str.,* as originally published by Mueller (1858).

Although *P. phylicifolia, P. gilesii* and *P.* sp. Evans Crown are closely related, gene flow is highly restricted, and the phenetic differences detected in this study suggest that there are likely genetically linked phenotypic traits that have been naturally selected for. The solution that most accurately represents the extent of genetic and morphological variation within this clade is to treat all three taxa as independently evolving metapopulation lineages (*sensu* de Queiroz 2007) that warrant species-level recognition. *Prostanthera* sp. Evans Crown is demonstrably genetically and morphologically distinct from *P. gilesii* and *P. phylicifolia* and is consequently described here as *P. volucris* R.P.O’Donnell to accommodate accessions from the Tarana region of New South Wales.

### Taxonomy

#### 1. Prostanthera phylicifolia

F.Muell. *Fragmenta Phytographiæ Australiæ* 1(1): 19 (1858).

##### Protologue citation

“In vertice rupestri montis McFarlane altitudine 4–5000’, nec non in rupibus secus rivulos districtus Maneroo.” = “At the summit of rocky mountain McFarlane from altitudes of 4–5000’, and also from cliffs or small rivulets in the district Maneroo.”

##### Lectotype (designated here)

Australia: Victoria: Eastern Highlands: “Mt M’Farlan”, *F. Mueller s. n.* (K000975463: left-hand side of sheet, right-hand sprig); isolectotypes: K000975463: left-hand side of sheet, left-hand sprig; residual syntypes: MEL43499 (right-hand side of sheet), MEL43500, K000975460 (Maneroo).

##### Notes

There are several specimens of *P. phylicifolia* that are considered to be collections made by Mueller (MEL43499: left-hand side, MEL43501, K000975461); however, the handwriting on these respective accessions does not match that of Mueller’s and is subsequently of uncertain authorship. As it is unclear whether these accessions were indeed collected and inspected by Mueller, they cannot be considered to have any nomenclatural standing and are not cited here. Specimen K000975462 is a collection from a locality described in Mueller’s handwriting as “Mitta Mitta”. This locality is not mentioned in Mueller’s protologue, and thus cannot be considered to have any nomenclatural standing.

Specimen K000975461 bears a label of uncertain authorship reading “var. velutina”, and K000975462 is similarly labelled with “v. velutina”, albeit in this instance written definitively in Mueller’s handwriting. This variety is not recognised in APNI (2021) or (IPNI 2021) and while Mueller’s protologue (1858, p. 19) describes individuals of *P. phylicifolia* as “glabra v. velutina” (i.e. glabrous or velutinous), neither of these qualifiers are separated by Mueller as named varieties. Therefore, the name *P. phylicifolia* var. *velutina* appears to be a manuscript name of no nomenclatural standing and has never been validly published. O’Donnell et al. (2021) found little genetic differentiation between individuals of *P. phylicifolia* that were predominantly glabrous and individuals that were densely hairy, suggesting that these character states are of dubious taxonomic utility. As most specimens of *P. phylicifolia* examined in this study were predominantly glabrous and some of the glabrous syntypes were well endowed with flowers and seemingly fruiting calyces, a representatively glabrous specimen was designated as the lectotype.

As O’Donnell et al. (2021) demonstrated that *P. phylicifolia s. str.* is restricted to southern New South Wales and Victoria, Mueller’s locality description of “Maneroo” must be considered to be an early spelling of Monaro (referring to the Monaro region), and not the rural locality of Maneroo located in Longreach, Queensland.

#### 2. Prostanthera volucris R.P.O’Donnell, *sp. nov.* (Figures 7, 8)

**Fig. 7.**
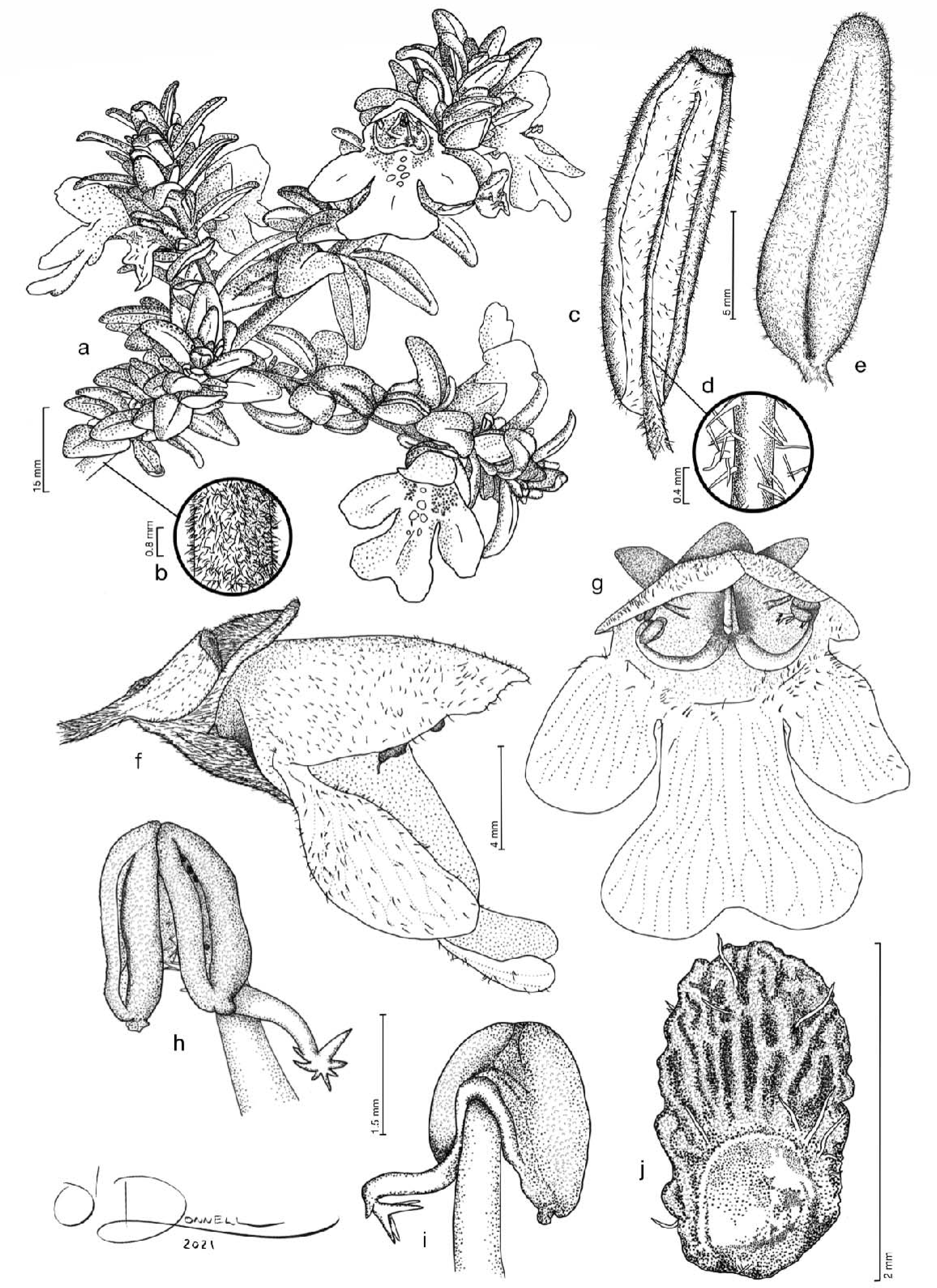
Illustration of *Prostanthera volucris* a. habit; b. detail of branch surface, showing retrorse trichomes; c. leaf surface, abaxial view; d. detail of abaxial leaf lamina surface, showing midrib and indumentum; e. leaf lamina surface, adaxial view; f. flower, lateral view, showing calyx, prophyll, corolla, anthers; g. flower, ventral view, showing corolla inner surface of lobes and tube, stamens, and style; h. stamen showing ventral view of anther locules, connective appendage and distal portion of staminal filament; i. stamen showing dorsal view of anther, connective appendage and distal portion of staminal filament; j. mericarp, ventral view, showing abscission scar. Illustration: R. P. O’Donnell

**Fig. 8.**
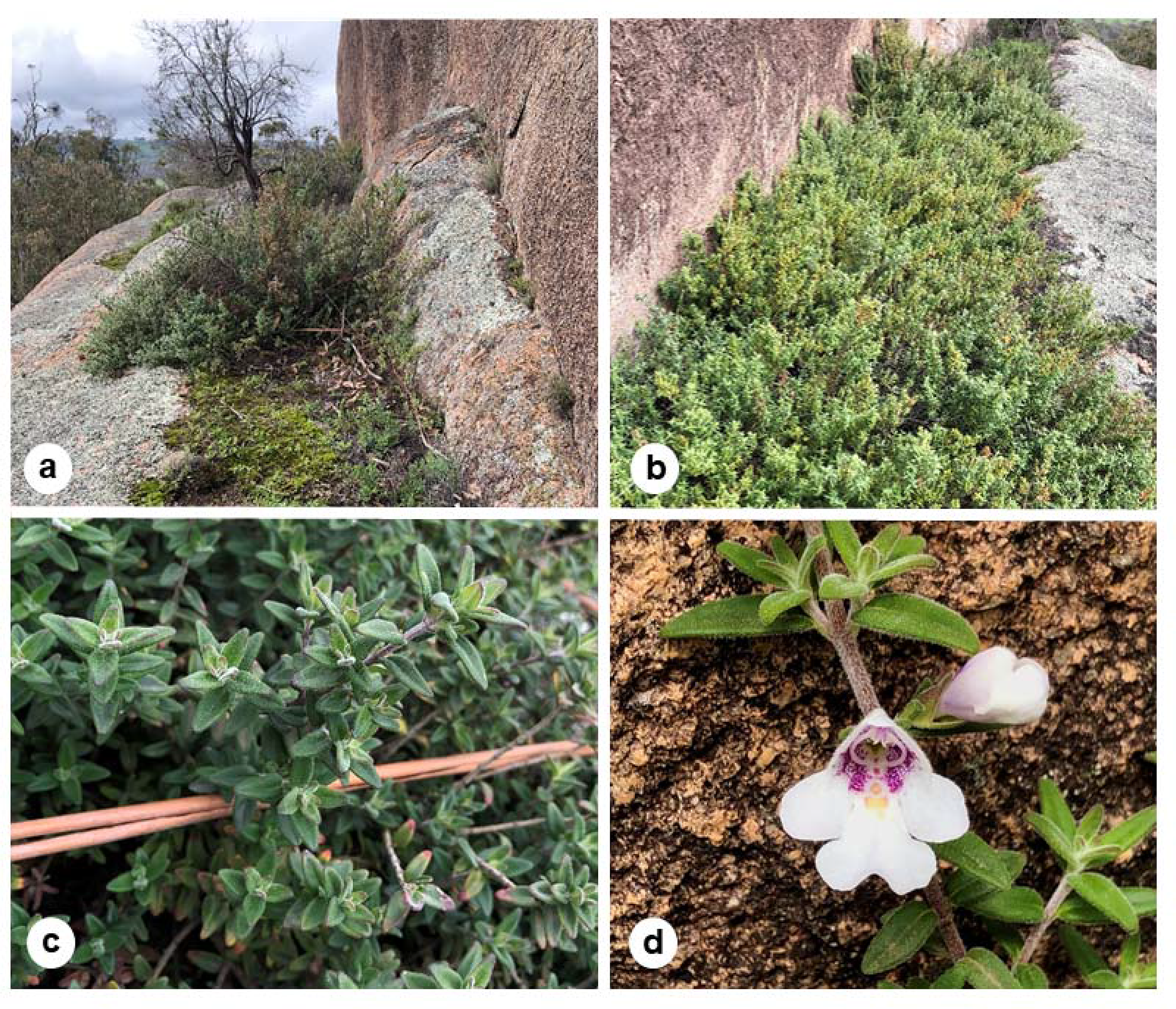
Photograph images of *Prostanthera volucris* a. habitat and associated vegetation; b. habit; c. habit, close up; d. flower and bud. Images: R. P. O’Donnell

##### Diagnosis

*Prostanthera volucris* can be distinguished from the morphologically similar *P. gilesii* by its larger prophylls, 3.8–9 mm long, 0.8–4.5 mm wide (vs. <4 mm long, <0.6 mm wide for *P. gilesii*), densely hairy branches, calyces and prophylls (up to 80 trichomes/mm^2^ vs. up to 40 trichomes/mm^2^ for *P. gilesii*), appressed to subappressed retrorse trichomes (vs. antrorse for *P. gilesii*) and densely hairy abaxial and adaxial lamina surfaces (vs. predominantly glabrous with occasional antrorse trichomes restricted to the abaxial surface midrib for *P. gilesii*). *Prostanthera volucris* can also be distinguished from *P. phylicifolia* and *P. gilesii* by its mericarps that are rugose and papillose with occasional long pilose trichomes (vs reticulate, not distinctly papillose and glabrous for *P. phylicifolia*; mature mericarps have never been observed for *P. gilesii*) (Table 4).

**Table 4.**
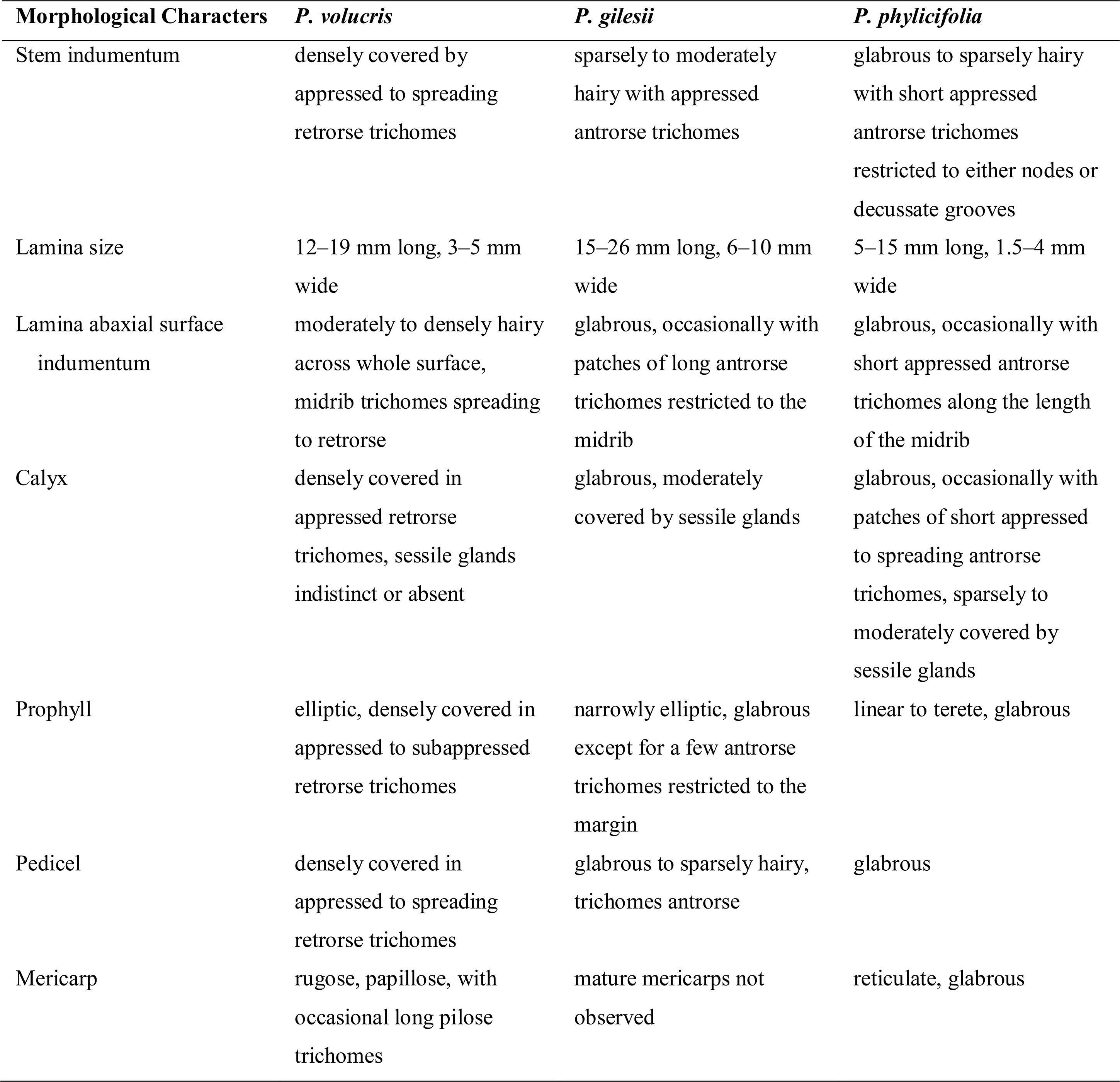
Selected morphological attributes separating *Prostanthera volucris*, *P. gilesii* and *P. phylicifolia*

##### Holotype

Australia: New South Wales: Central Tablelands: Evans Crown Nature Reserve, c. 2.8 km SE of Tarana township, *G.M. Taseski 853, 28 October 2018* (NSW1055966: voucher; NSW1057497: spirit); isotypes: BRI, CANB, K, MEL, MO, NE110628, P, UNSW.

##### Informal phrase names

*Prostanthera* sp. Evans Crown (G.M.Taseski NSW1055966) (O’Donnell et al. 2021) and *P.* sp. nov. aff. *gilesii.*

Compact, erect shrub up to 80 cm high, with populations forming tight mats. *Branches ±* terete, occasionally quadrangular early in development, becoming terete with age, densely hairy [up to c. 80 trichomes/mm^2^]; trichomes 0.1–0.4 mm long, appressed to subappressed, retrorse, straight to slightly curled, white; sessile glands indistinct or absent (obscured by indumentum). *Leaves* velutinous, appearing silver–light green, slightly paler on abaxial surface, occasionally becoming red after prolonged exposure to strong sunlight, not aromatic when touched or crushed; *petiole* 0.88–1.4 mm long, densely hairy [20–48 trichomes/mm^2^]; trichomes 0.2–0.8 mm long, appressed to subappressed, retrorse, straight to slightly curled, white; *lamina* narrowly ovate to elliptic, 12–19 mm long, 3–5 mm wide, length to width ratio 2.6–5.3, length of maximum width from base to total lamina length ratio 0.1–0.4; moderately to densely hairy on abaxial surface, particularly on midrib [up to c. 40 trichomes/mm^2^], trichomes 0.2–0.4 mm long, spreading to erect, occasionally retrorse along midrib, straight to slightly curled, white, glands indistinct; moderately to densely hairy on adaxial surface [up to c. 30 trichomes/mm^2^], trichomes 0.1–0.3 mm long, spreading to erect, straight to slightly curled, white, glands indistinct; base obtuse, occasionally appearing attenuate because margin more involute towards base; margin recurved; apex obtuse; venation indistinct, midrib slightly raised on abaxial surface. *Inflorescence* a frondose botryoidal conflorescence, uniflorescence monadic; 4–8-flowered (per conflorescence)*. Podium* a_1_ axis 0.9–2.8 mm long, densely hairy [30–60(–88) trichomes/mm^2^]; trichomes 0.1–0.2 mm long, appressed to subappressed, retrorse, straight to slightly curled, white; glands indistinct or absent (obscured by indumentum); anthopodium absent or indistinct. *Pherophylls* not seen. *Prophylls ±* persistent, inserted near base of calyx, opposite, narrow ovate to narrowly elliptic, 3.8–9 mm long, 0.8–4.5 mm wide, length to width ratio 1.8–5.6, length of maximum width from base to total lamina length ratio 0.3–0.9, densely hairy [up to 56 trichomes/mm^2^]; trichomes 0.1–0.2 mm long, appressed to subappressed, retrorse, straight to slightly curled, white; glands indistinct or absent (obscured by indumentum); base slightly attenuate; margin entire; apex obtuse; venation indistinct. *Calyx tube* 1.5–2.1 mm long; *abaxial lobe* ovate to broadly ovate, 2–4 mm long, 2–4 mm wide at base, apex rounded to retuse; *adaxial lobe* ovate to elliptic, 3–5.3 mm long (adaxial lobe length to abaxial lobe length ratio c. 1.3), 2.1–4.2 mm wide at base, length to width ratio 1.2–1.4, apex *±* rounded; outer surface densely hairy [up to 64 trichomes/mm^2^], trichomes 0.2–0.35 mm long, appressed to subappressed, retrorse, straight to slightly curled, white, glands indistinct or absent (obscured by indumentum); inner surface of tube glabrous, lobes moderately to densely hairy near margin and apex, trichomes appressed to subappressed, spreading to occasionally antrorse, straight to slightly curled, white; light green, occasionally darkening to dark mauve with sun exposure. *Corolla* 14–18 mm long; *tube* 3–5 mm long; *abaxial median lobe* broadly spathulate, 7–9.5mm long, 4–7mm wide (below distal lobing), length to width ratio 1.4–1.8, apex slightly irregular and rounded, bilobed (sinus 2–2.3 mm long, 2.5–3 mm wide distally); *lateral lobes* oblong to slightly elliptic, 5.4–5.7 mm long, 3.1–3.8 mm wide, length to width ratio 1.5–1.8, apex rounded, slightly irregular; *adaxial median lobe-pair* broad to depressed ovate, 6–7 mm long, 9–9.5 mm wide, length to width ratio 0.6–0.8, apex rounded, irregular, bilobed (sinus 0.6–0.8 mm long, 0.9–1.6 mm wide, median margin of lobes occasionally overlapping slightly); outer surface moderately hairy, particularly on lobes [8–24 trichomes/mm^2^]; trichomes 0.1–0.4 mm long, erect to spreading along tube before becoming appressed to subappressed and antrorse on lobes, sparsely glandular; inner surface *±* glabrous, lobes sparsely hairy to moderately hairy at the tube opening rim and sinuses between lobes, trichomes 0.1–0.2 mm long, crinkled; white, with purple to dark mauve speckled markings on the inner surface of the tube, with pale orange to yellow markings on base of abaxial median lobe. *Stamens* inserted 2.8–4.1 mm above base of corolla; *filaments* 3.1–5.4 mm long, white, often with mauve tinge; *anthers* 1.4–1.8 mm long, minutely papillose with an acumen at the base of each lobe and trichomes between lobes, trichomes 0.1–0.2 mm long, white; connective appendage 0.8–1.5 mm long, with a few narrowly triangular trichomes 0.1–0.35 mm long, white; dark mauve maturing to yellow-brown. *Disc* 0.5–1 mm long. *Pistil* 7.5–9.5 mm long; *ovary* cylindrical obovoid, 0.9–1 mm long, at base 0.9–1 mm diameter, lobes 0.5–0.65 mm long; *style* 7.5–8 mm long; *stigma lobes* 0.3–0.4 mm long. *Fruiting calyx* not strongly accrescent, only slightly enlarged, abaxial lobe clasping to conceal developing mericarps, adaxial lobe not strongly reflexed. *Mature mericarps* 1.8–2.1 mm long, 1–1.2 mm wide, rugose, minutely papillose, with a few spreading trichomes; trichomes 0.2–0.5 mm long, white.

##### Distribution

Known only from a single granitic tor in the Evans Crown Nature Reserve, SE of Tarana, New South Wales, Australia (Figure 1). This location is situated within the Central Tablelands Botanical Division (*South Eastern Highlands Biogeographic Region*).

##### Habitat

This species grows on exposed granite formations at c. 1,000–1,020 m altitude, along drainage crevices in shallow, skeletal humic soils with *Cyphanthera albicans* and *Cheilanthes sieberi* nearby.

##### Etymology

The specific epithet ‘*volucris*’ (‘winged’ or ‘winged creature’) refers to this species’ substantial and densely hairy prophylls. Inserted at the base of the calyx, the large prophylls give the calyx the appearance of being ‘winged’.

##### Ecology and conservation

While the first collector of this species considered it “locally abundant in rock crevices” (McKee 7043, NSW 237164), only one population has been located in subsequent searches. Leaf material was sampled from a herbarium voucher specimen (Rodd 11009, NSW 856887) that was collected from a population with coordinates that do not match the known, densely sampled population. Unfortunately, DNA extraction for that sample failed to yield sufficient DNA material for sequencing. The coordinates indicated on this voucher should be surveyed as a priority. More intensive surveys of the Evans Crown Nature Reserve and vegetation islands in the region are needed to look for additional populations. The Tarana region experienced extensive land clearing following European settlement in the area in the 1860s and the Evans Crown area was leased for grazing from 1880–1972 (NPWS 2009). A small granite quarry was also in operation in the reserve in the early 1940s (NPWS 2009), and as *P. volucris* is known to occur on granite outcrops, it is probable that mining operations resulted in a reduction of suitable habitat options. It is likely that *P. volucris* has suffered substantial declines in distribution following European settlement. It is unknown whether *P. volucris* is subject to herbivory; however, sheep and goats have been known to access the reserve from neighbouring properties and there are currently no feral animal control programs implemented for the reserve (NPWS 2009). The reserve is a popular attraction for hikers and rock climbers, and as the species occurs on a geologically striking outcrop, human activity is a likely threat to this population. Upon revisiting the site following the severe drought conditions of 2019–2020, the population appeared to be severely affected by heat and drought stress and had declined substantially in condition (Taseski pers. obs. 2020). It is probable that projected prolonged drought conditions and heightened temperatures will adversely affect this population in a severe manner, particularly on account of its highly exposed habitat. As *P. volucris* grows in tight, tangled mats, it is difficult to determine the limits of individual plants. Genomic relationship matrices (Online Resource 7) indicate that some samples represent ramets from a clonal parent, while others represent multiple genetically distinct individuals. Seedlings have been observed in recent surveys and mature individuals have been observed to set seed, however, it is unknown what proportion of seed set is viable. As *P. volucris* is known to occur along drainage crevices, it is likely that seeds are dispersed primarily by water runoff. The pollinators of *P. volucris* are unknown, however, the combination of floral characteristics that it exhibits corresponds with a floral type visited primarily by bees, although not excluding other insects such as flies (Wilson et al. 2017). Studies of maximum foraging ranges in Australian bees found a typical maximum foraging range of approximately 700 m (Smith et al. 2017), which suggests that pollen is unlikely to travel further than this distance.

Given the horticultural potential of *P. volucris,* this species could be a target of increased collecting pressure. Attempts are underway to develop an *ex situ* collection with botanic gardens and native plant wholesalers. The known distribution of this species is restricted such that the area of occupancy and extent of occurrence are not greater than 4 km^2^. As this species is known from only one highly restricted population in a <1 km^2^ area, and has an estimated population of <250 mature individuals, we suggest that this species satisfies the criteria to be considered Critically Endangered under the *New South Wales Biodiversity Conservation Act* (2016), the *Environment Protection and Biodiversity Conservation Act* (1999b) and the *IUCN Red List* criteria thresholds (2019).

##### Notes

While mericarp characters were not scored for morphological analyses, differences in mericarp morphology appear to be taxonomically informative between *P. volucris* and *P. phylicifolia* (rugose and distinctly papillate in *P. volucris* vs. reticulate and not distinctly papillose in *P. phylicifolia*). Scant attention has been paid to mericarp surface ornamentation and sculpting in recent descriptions of *Prostanthera,* with the exception of Williams et al. (2006) in their treatment of the *P. spinosa* complex and Guerin’s (2005) study of mericarp morphology in the Westringieae. The findings outlined here and by Guerin (2005) and Williams et al. (2006) highlight that mericarp morphology may be taxonomically informative in *Prostanthera*. Further examination of mericarp morphology across the genus is warranted and may provide additional characters for distinguishing between closely related species.

##### Other specimens examined

AUSTRALIA: NEW SOUTH WALES: CENTRAL TABLELANDS: (*South Eastern Highlands*): Evans Crown Nature Reserve: *H.S. McKee 7043*, 10 Jan. 1960 (NSW237164); *A.N. Rodd 11009*, 9 Mar. 2002 (NSW856887); *R.P. O’Donnell & G.M. Taseski 28,* 7 Apr. 2020 (NSW1100357); *R.P. O’Donnell & G.M. Taseski 29,* 7 Apr. 2020 (NSW1100369); *R.P. O’Donnell & G.M. Taseski 30,* 7 Apr. 2020 (NSW1100379); *R.P. O’Donnell & T.C. Wilson 55*, 4 Oct. 2020 (NSW1100402); *R.P. O’Donnell & T.C. Wilson 56*, 4 Oct. 2020 (NSW1100403).

## Supporting information

Online Resources

## Acknowledgements

The National Herbarium of New South Wales and the N.C.W. Beadle Herbarium provided access to their *Prostanthera* collections, and the Research Centre for Ecosystem Resilience (Royal Botanic Gardens and Domain Trust) provided us access to their DArTseq dataset of *P. densa* and *P. marifolia* for co-analysis. We thank Richard Medd for collections and discussion about *P. gilesii* and *P. phylicifolia*; Neville Walsh (MEL) for silica gel material from Victorian populations of *P. phylicifolia*; Danielle Smith and Helen Kennedy (NE) for preparation of leaf samples for genetic analysis; Lesley Elkan (NSW) for constructive feedback on illustrations; Ben Correy and Shane Smith (NSW National Parks and Wildlife Service) for correspondence and logistics pertaining to Evans Crown Nature Reserve; Markus Buchhorn for access to and assistance on his property, and Stephen Hopper (UWA) and Katharina Nargar (CNS) for constructive comments on the manuscript. Fieldwork was conducted under New South Wales OEH Scientific Licence SL100305.

## Statements and Declarations

### Author contributions

All authors contributed to the conception and design of this study. Data collection and analysis were performed by Ryan P. O’Donnell. The first draft of the manuscript was written by Ryan P. O’Donnell and all authors contributed comments and revisions to subsequent versions. All authors read and approved the final manuscript.

### Data availability

The datasets generated and analysed in this study are available from the corresponding author upon reasonable request.

