## Supplementary material for "Molecular and morphological analyses support recognition of *Prostanthera volucris* (Lamiaceae), a new species from the Central Tablelands of New South Wales": Online Resources

### Online Resource 1 Samples of *Prostanthera* that were sequenced and co-analysed to produce DArTseq SNP data, arranged alphabetically by species. NP = National Park; NR = Nature Reserve; SF = State Forest; SCA = State Conservation Area; N.S.W. = New South Wales; Vic. = Victoria. * = newly sequenced for this study; ^ = technical duplicates; ° = ReCER samples; § = removed by filtering processes. A = primary SNP dataset; B = secondary phylogenetic dataset; ! = individuals retained for genetic diversity calculations following removal of putative clones

| **Species** | **Locality** | **OTU Code** | **Voucher** | **Herbarium and accession no.** | **A** | **B** |
| --- | --- | --- | --- | --- | --- | --- |
| *P. densa*  A.A. Ham. | Gaan Gaan Hill, N.S.W. | - | Wood *s.n.*° | NSW1040587 | - | 🗸 |
|  | Beecroft Peninsula, N.S.W | - | Conn 2571° | NSW203752 | - | 🗸 |
| *P. gilesii*  G.W.Althofer *ex* B.J.Conn & T.C.Wilson | Towac Creek, Mount Canobolas SCA, N.S.W. | HZ1 | Zimmer 1 | - | 🗸^!^ | 🗸 |
|  |  | HZ1.1 | Zimmer 1.1^ | - | 🗸^§^ | - |
|  |  | HZ2 | Zimmer 2 | - | 🗸 | - |
|  |  | HZ3 | Zimmer 3 | - | 🗸 | - |
|  |  | HZ4 | Zimmer 4 | - | 🗸 | - |
|  |  | HZ75 | Zimmer 75 | - | 🗸 | - |
|  |  | HZ76 | Zimmer 76 | - | 🗸 | - |
|  |  | HZ77 | Zimmer 77 | - | 🗸 | - |
|  |  | HZ79 | Zimmer 79 | - | 🗸^!^ | - |
|  |  | HZ81 | Zimmer 81 | - | 🗸 | - |
|  |  | HZ82 | Zimmer 82 | - | 🗸 | - |
|  |  | HZ83 | Zimmer 83 | - | 🗸 | - |
|  |  | HZ90 | Zimmer 90 | - | 🗸 | - |
|  |  | HZ91 | Zimmer 91 | - | 🗸 | - |
|  |  | HZ92 | Zimmer 92 | - | 🗸 | - |
|  |  | HZ93 | Zimmer 93 | - | 🗸 | - |
|  |  | HZ93.1 | Zimmer 93.1^ | - | 🗸^§^ | - |
|  |  | HZ94 | Zimmer 94 | - | 🗸 | - |
|  |  | HZ95 | Zimmer 95 | - | 🗸 | - |
|  |  | HZ96 | Zimmer 96 | - | 🗸 | - |
|  |  | HZ97 | Zimmer 97 | - | 🗸 | - |
|  |  | HZ101 | Zimmer 101 | - | 🗸 | - |
|  |  | HZ107 | Zimmer 107 | - | 🗸 | - |
|  |  | HZ109 | Zimmer 109 | - | 🗸 | - |
|  |  | HZ111 | Zimmer 111 | - | 🗸 | - |
|  |  | HZ113 | Zimmer 113 | - | 🗸 | - |
|  |  | HZ115 | Zimmer 115 | - | 🗸 | - |
|  |  | JJB3602a | Bruhl 3602a | NE108967 | 🗸^!^ | 🗸 |
|  |  | JJB3602aa | Bruhl 3602aa | “ | 🗸 | - |
|  |  | JJB3602b | Bruhl 3602b | “ | 🗸 | - |
|  |  | JJB3602bb | Bruhl 3602bb | “ | 🗸 | - |
|  |  | JJB3602c | Bruhl 3602c | “ | 🗸 | - |
|  |  | JJB3602cc | Bruhl 3602cc | “ | 🗸 | - |
|  |  | JJB3602d | Bruhl 3602d | “ | 🗸 | - |
|  |  | JJB3602dd | Bruhl 3602dd | “ | 🗸 | - |
|  |  | JJB3602e | Bruhl 3602e | “ | 🗸 | - |
|  |  | JJB3602ee | Bruhl 3602ee | “ | 🗸^§^ | - |
|  |  | JJB3602ee.1 | Bruhl 3602ee.1^ | “ | 🗸 | - |
|  |  | JJB3602f | Bruhl 3602f | “ | 🗸 | - |
|  |  | JJB3602g | Bruhl 3602g | “ | 🗸 | - |
|  |  | JJB3602h | Bruhl 3602h | “ | 🗸 | - |
|  |  | JJB3602i | Bruhl 3602i | “ | 🗸 | - |
|  |  | JJB3602j | Bruhl 3602j | “ | 🗸 | - |
|  |  | JJB3602k | Bruhl 3602k | “ | 🗸 | - |
|  |  | JJB3602l | Bruhl 3602l | “ | 🗸 | - |
|  |  | JJB3602m | Bruhl 3602m | “ | 🗸 | - |
|  |  | JJB3602n | Bruhl 3602n | “ | 🗸 | - |
|  |  | JJB3602o | Bruhl 3602o | “ | 🗸 | - |
|  |  | JJB3602p | Bruhl 3602p | “ | 🗸 | - |
|  |  | JJB3602q | Bruhl 3602q | “ | 🗸 | - |
|  |  | JJB3602r | Bruhl 3602r | “ | 🗸 | - |
|  |  | JJB3602s | Bruhl 3602s | “ | 🗸 | - |
|  |  | JJB3602t | Bruhl 3602t | “ | 🗸 | - |
|  |  | JJB3602u | Bruhl 3602u | “ | 🗸 | - |
|  |  | JJB3602v | Bruhl 3602v | “ | 🗸 | - |
|  |  | JJB3602w | Bruhl 3602w | “ | 🗸 | - |
|  |  | JJB3602x | Bruhl 3602x | “ | 🗸 | - |
|  |  | JJB3602y | Bruhl 3602y | “ | 🗸 | - |
|  |  | JJB3602z | Bruhl 3602z | “ | 🗸 | - |
|  |  | JJBsnf1 | Bruhl snf1 | - | 🗸 | - |
|  |  | JJBsnf2 | Bruhl snf2 | - | 🗸 | - |
|  |  | JJBsnf3 | Bruhl snf3 | - | 🗸 | - |
|  |  | JJBsng | Bruhl sng | - | 🗸 | - |
|  |  | TCW581 | Wilson 581 | - | 🗸 | - |
|  | The Walls, Mount Canobolas SCA, N.S.W. | TCW577 | Wilson 577 | NSW998442 | 🗸^!^ | 🗸 |
|  |  | RWM7111 | Medd 7111* | - | 🗸^!^ | 🗸 |
| *P. granitica*  Maiden & Betche | Warrumbungles NP, N.S.W. | - | Tuft 162 (1)° | NSW1057815 | - | 🗸 |
|  |  | - | Tuft 162 (2)* | NSW844192 | - | 🗸 |
| *P. marifolia*  R.Br. | Seaforth, N.S.W. | - | Conn 5676° | NSW841013 | - | 🗸 |
|  | Wakehurst, N.S.W. | - | McVickar *s.n.*° | NSW1038335 | - | 🗸 |
|  | Manly, N.S.W. | - | Wilson 578° | NSW1057839 | - | 🗸 |
| *P. phylicifolia*  F.Muell. | Deua NP, N.S.W. | NSW1057814 | Liney 2039° | NSW1057814 | 🗸^§^ | 🗸 |
|  | Kosciuszko NP, N.S.W. | RWM2170961a | Medd 2170961a | NE109821 | 🗸^!^ | 🗸 |
|  |  | RWM2170961b | Medd 2170961b | “ | 🗸^!^ | - |
|  |  | RWM2170962a | Medd 2170962a | “ | 🗸^!^ | 🗸 |
|  |  | RWM2170962b | Medd 2170962b | “ | 🗸^!^ | - |
|  | Dangelong NR, N.S.W. | RWM217099a1 | Medd 217099a1 | NE109826 | 🗸^!^ | 🗸 |
|  |  | RWM217099b1 | Medd 217099b1 | “ | 🗸^!^ | 🗸 |
|  |  | RWM217099c1 | Medd 217099c1 | “ | 🗸^!^ | - |
|  |  | RWM217099d1 | Medd 217099d1 | “ | 🗸^!^ | - |
|  |  | RWM217099e1 | Medd 217099e1 | “ | 🗸^!^ | - |
|  | Adaminaby, N.S.W. | RWM217100a1 | Medd 217100a1 | NE109825 | 🗸^!^ | 🗸 |
|  |  | RWM217100b1 | Medd 217100b1 | “ | 🗸^!^ | 🗸 |
|  |  | RWM217100c1 | Medd 217100c1 | “ | 🗸^!^ | - |
|  |  | RWM217100d1 | Medd 217100d1 | “ | 🗸^!^ | - |
|  |  | RWM217100e1 | Medd 217100e1 | “ | 🗸^!^ | - |
|  | Nullica SF, N.S.W. | GPP219 | Phillips 219* | NSW992045 | 🗸^§^ | 🗸 |
|  |  | MP9561 | Parris 9561* | NSW456970 | 🗸^§^ | 🗸^§^ |
|  | Cobrunga, Vic. | NGW8979 | Walsh 8979* | MEL2470086 | 🗸^!^ | 🗸 |
|  |  | NGW9013 | Walsh 9013* | MEL2470120 | 🗸^!^ | 🗸 |
|  | Gelantipy, Vic. | ROM984 | Makinson 984* | NSW518048 | 🗸^§^ | 🗸^§^ |
|  | Tinderry, N.S.W. | RPO61 | O'Donnell 61* | NSW100407 | 🗸^§^ | 🗸 |
|  |  | RPO62 | O'Donnell 62* | NSW1100408 | 🗸^!^ | 🗸 |
| *P. scutellarioides* (R.Br.) Briq. | Castlereagh NR N.S.W. | - | Wilson 215° | NSW799702 | - | 🗸 |
|  | Myall River, N.W.W. | - | Bell *s.n.** | NSW847837 | - | 🗸^§^ |
| *P.* sp. Evans Crown | Evans Crown NR, N.S.W. | GMT853 | Taseski 853 | NSW1055966 | 🗸^§^ | - |
|  |  | GMT853a | Taseski 853a | “ | 🗸 | - |
|  |  | GMT853b | Taseski 853b | “ | 🗸 | - |
|  |  | GMT853c | Taseski 853c | “ | 🗸 | - |
|  |  | GMT853d | Taseski 853d | “ | 🗸^!^ | - |
|  |  | GMT853e | Taseski 853e | “ | 🗸^!^ | 🗸 |
|  |  | GMT853f | Taseski 853f | “ | 🗸 | - |
|  |  | RPO28a | O'Donnell 28a* | NSW1100357 | 🗸^!^ | 🗸 |
|  |  | RPO28b | O'Donnell 28b* | NSW1100358 | 🗸 | - |
|  |  | RPO28c | O'Donnell 28c* | NSW1100359 | 🗸 | - |
|  |  | RPO28d | O'Donnell 28d* | NSW1100360 | 🗸 | - |
|  |  | RPO28e | O'Donnell 28e* | NSW1100361 | 🗸 | - |
|  |  | RPO28f | O'Donnell 28f* | NSW1100362 | 🗸 | - |
|  |  | RPO28g | O'Donnell 28g* | NSW1100363 | 🗸 | - |
|  |  | RPO28i | O'Donnell 28i* | NSW1100365 | 🗸 | - |
|  |  | RPO28j | O'Donnell 28j* | NSW1100366 | 🗸 | - |
|  |  | RPO29a | O'Donnell 29a* | NSW1100369 | 🗸^!^ | 🗸 |
|  |  | RPO29b | O'Donnell 29b* | NSW1100370 | 🗸 | - |
|  |  | RPO29d | O'Donnell 29d* | NSW1100412 | 🗸 | - |
|  |  | RPO29e | O'Donnell 29e* | NSW1100373 | 🗸^!^ | - |
|  |  | RPO29f | O'Donnell 29f* | NSW1100374 | 🗸 | - |
|  |  | RPO29g | O'Donnell 29g* | NSW1100375 | 🗸 | - |
|  |  | RPO29h1 | O'Donnell 29h1* | NSW1100376 | 🗸 | - |
|  |  | RPO29h2 | O'Donnell 29h2* | “ | 🗸 | - |
|  |  | RPO29i | O'Donnell 29i* | NSW1100377 | 🗸 | - |
|  |  | RPO29j | O'Donnell 29j* | NSW1100378 | 🗸 | - |
|  |  | RPO30b | O'Donnell 30b* | NSW1100380 | 🗸^!^ | - |
|  |  | RPO30c | O'Donnell 30c* | NSW1100381 | 🗸^§^ | - |
|  |  | RPO30d | O'Donnell 30d* | NSW1100382 | 🗸 | - |
|  |  | RPO30e | O'Donnell 30e* | NSW1100383 | 🗸 | - |
|  |  | RPO30f | O'Donnell 30f* | NSW1100384 | 🗸 | - |
|  |  | RPO30g | O'Donnell 30g* | NSW1100385 | 🗸 | - |
|  |  | RPO30h | O'Donnell 30h* | NSW1100386 | 🗸 | - |
|  |  | RPO30i | O'Donnell 30i* | NSW1100387 | 🗸 | - |
|  |  | RPO30j | O'Donnell 30j* | NSW1100388 | 🗸 | - |
|  |  | RPO55 | O'Donnell 55* | NSW1100402 | 🗸^§^ | 🗸 |
|  |  | RPO56 | O'Donnell 56* | NSW1100403 | 🗸^!^ | - |

### Online Resource 2 Voucher specimens and OTU codes of *Prostanthera* measured for multivariate phenetic analysis in this study. NP = National Park; NR = Nature Reserve; SF = State Forest; SCA = State Conservation Area; A.C.T. = Australian Capital Territory; N.S.W. = New South Wales; Vic. = Victoria

| **Taxon** | **OTU code** | **Locality** | **Voucher** | **Herbarium and accession no.** |
| --- | --- | --- | --- | --- |
| *P.* sp. Evans Crown | EC1 | Evans Crown NR, N.S.W. | Rodd 11009 | NSW856887 |
|  | EC2 | Evans Crown NR, N.S.W. | McKee 7043 | NSW237164 |
|  | EC3 | Evans Crown NR, N.S.W. | Taseski 853 | NSW1055966 |
|  | EC4 | Evans Crown NR, N.S.W. | O'Donnell 55 | NSW1100402 |
|  | EC5 | Evans Crown NR, N.S.W. | O'Donnell 28 | NSW1100357 |
| *P. gilesii* | gil1 | Mt Canobolas SCA, N.S.W. | Bruhl 3601b | NE108967 |
|  | gil2 | Mt Canobolas SCA, N.S.W. | Bruhl 3619 | NE109460 |
|  | gil3 | Mt Canobolas SCA, N.S.W. | Bruhl 3601d | NE108967 |
|  | gil4 | Mt Canobolas SCA, N.S.W. | Giles s.n. | NSW128315 |
|  | gil5 | Mt Canobolas SCA, N.S.W. | Giles s.n. | NSW128313 |
| *P. phylicifolia* | phyCobb | Mt Cobberas #2, Vic. | Walsh 2039 | NSW243340 |
|  | phyWul | Wulgulmerang, Vic. | Melville 3036 | NSW1051719 |
|  | phyCoop | Mt Coopracambra, Vic. | Albrecht 3684 | NSW1051502 |
|  | phyAda | Adaminaby, N.S.W. | Medd 217100a | NE109830 |
|  | phyKNP | Kosciuszko NP, N.S.W. | Medd 217096b | NE109822 |
|  | phyDang | Dangelong NR, N.S.W. | Medd 217099b | NE109826 |
|  | phyDeua | Big Badja Hill, Deua NP, N.S.W. | Liney 2039 | NSW704841 |
|  | phyNam | Namadgi NP, A.C.T. | Taws 315 | NSW624610 |
|  | phyTur | Tuross River, N.S.W. | Briggs s.n. | NSW275953 |
|  | phyTin | Tinderry Ranges, N.S.W. | O'Donnell 61 | NSW1100407 |

### Online Resource 3 Morphological characters used in multivariate phenetic analysis of selected specimens of *P. gilesii, P. phylicifolia* and *P.* sp. Evans Crown

| **Character no.** | **Character (units)** | **Character no.** | **Character (units)** |
| --- | --- | --- | --- |
| 1 | Branch hair direction:  trichomes absent (0); trichomes antrorse (1); trichomes retrorse (2); trichomes spreading (3) | 16 | Prophyll length (mm) |
| 2 | Branch hair density (trichomes/mm^2^) | 17 | Prophyll width (mm) |
| 3 | Branch hair length (mm) | 18 | Prophyll L:W ratio |
| 4 | Leaf petiole hair direction: trichomes absent (0); trichomes antrorse (1); trichomes retrorse (2); trichomes spreading (3) | 19 | Prophyll trichome density (trichomes/mm^2^) |
| 5 | Leaf lamina length (mm) | 20 | Prophyll hair direction: trichomes absent (0); trichomes antrorse (1); trichomes retrorse (2); trichomes spreading (3) |
| 6 | Leaf lamina width (mm) | 21 | Calyx outer surface hair direction: trichomes absent (0); trichomes antrorse (1); trichomes retrorse (2); trichomes spreading (3) |
| 7 | Leaf lamina L:W ratio | 22 | Calyx outer surface hair density (trichomes/mm^2^) |
| 8 | Leaf lamina abaxial surface hair density (trichomes/mm^2^) | 23 | Calyx abaxial lobe length (mm) |
| 9 | Leaf lamina abaxial surface hair length (mm) | 24 | Calyx abaxial lobe width (mm) |
| 10 | Leaf lamina abaxial surface midrib hair direction: trichomes absent (0); trichomes antrorse (1); trichomes retrorse (2); trichomes spreading (3) | 25 | Calyx abaxial lobe L:W ratio |
| 11 | Leaf lamina adaxial surface hair density (trichomes/mm^2^) | 26 | Calyx adaxial lobe length (mm) |
| 12 | Leaf lamina adaxial surface hair length (mm) | 27 | Calyx adaxial lobe width (mm) |
| 13 | a_1_ axis length (mm) | 28 | Calyx adaxial lobe L:W ratio |
| 14 | a_1_ axis hair density (trichomes/mm^2^) | 29 | Calyx abaxial:adaxial lobe length ratio |
| 15 | a_1_ axis hair direction:  trichomes absent (0); trichomes antrorse (1); trichomes retrorse (2); trichomes spreading (3) |  |  |

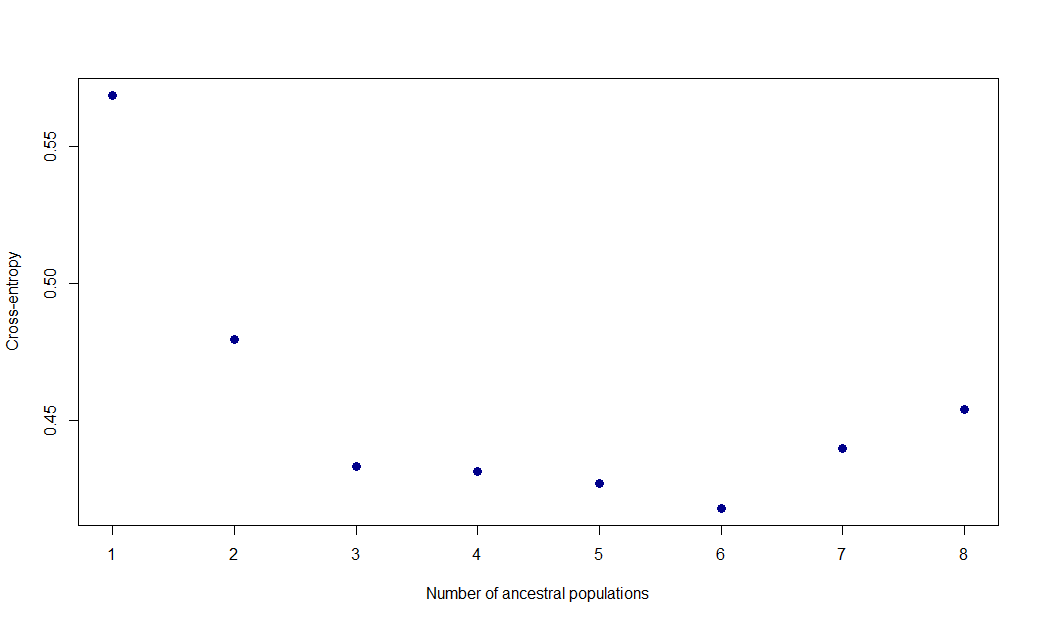

### Online Resource 4 Cross-entropy value plots with a stabilisation point of *K* = 3 indicating three probable ancestral populations represented in the data

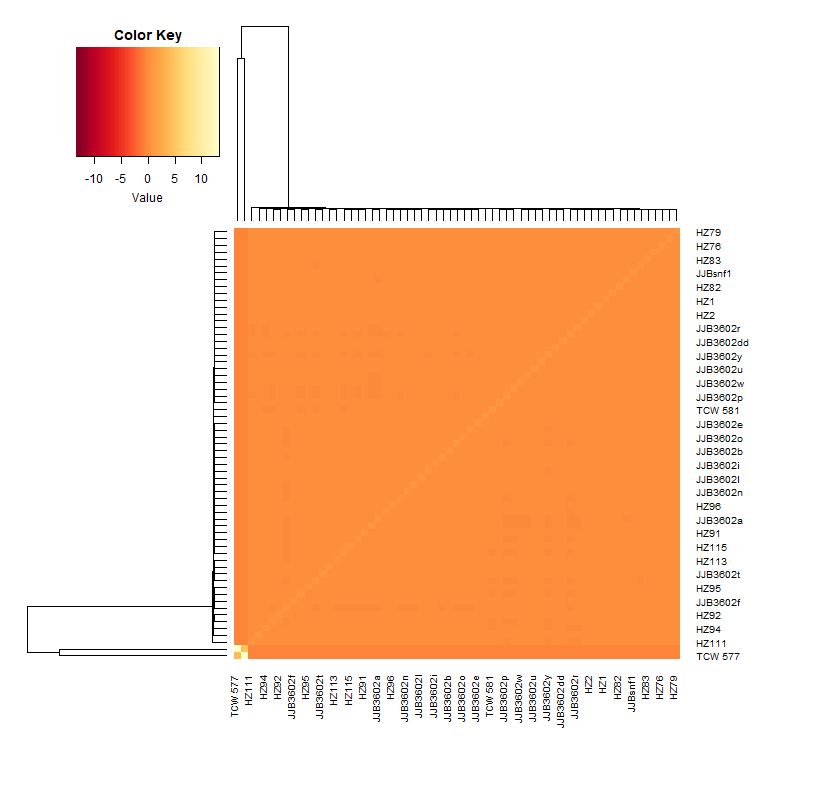

### Online Resource 5 Genomic relationship matrices for all sampled individuals of *P. gilesii*. Lighter colours correspond with individuals that are more closely related, with the lightest values at the central diagonal representing the relationship between each specimen and itself. Sample codes follow those outlined in Online Resource 1

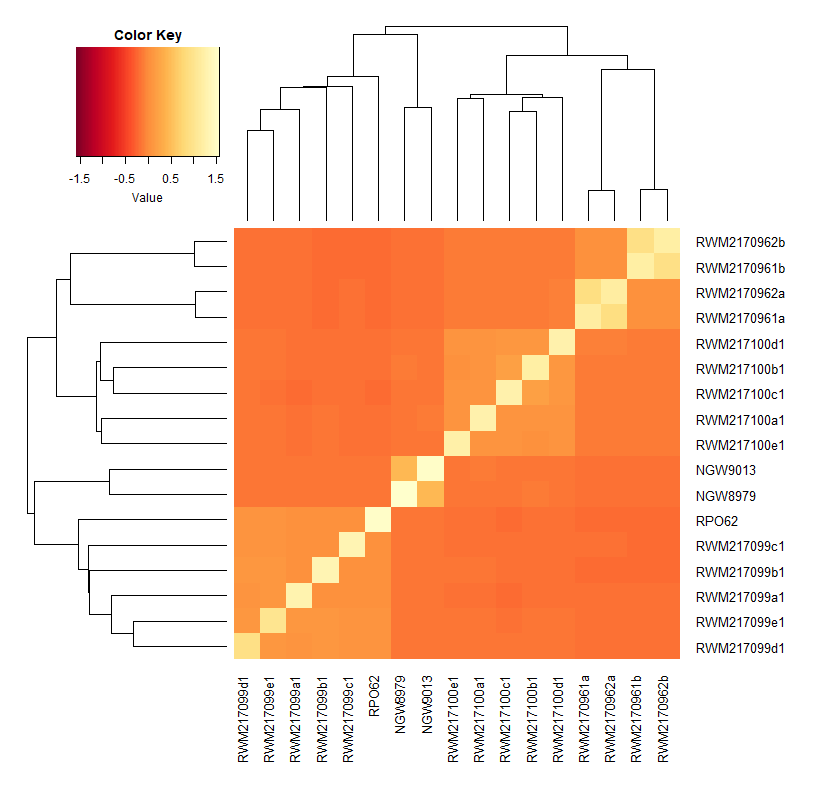

### Online Resource 6 Genomic relationship matrices for all sampled individuals of *P. phylicifolia*. Lighter colours correspond with individuals that are more closely related, with the lightest values at the central diagonal representing the relationship between each specimen and itself. Sample codes follow those outlined in Online Resource 1

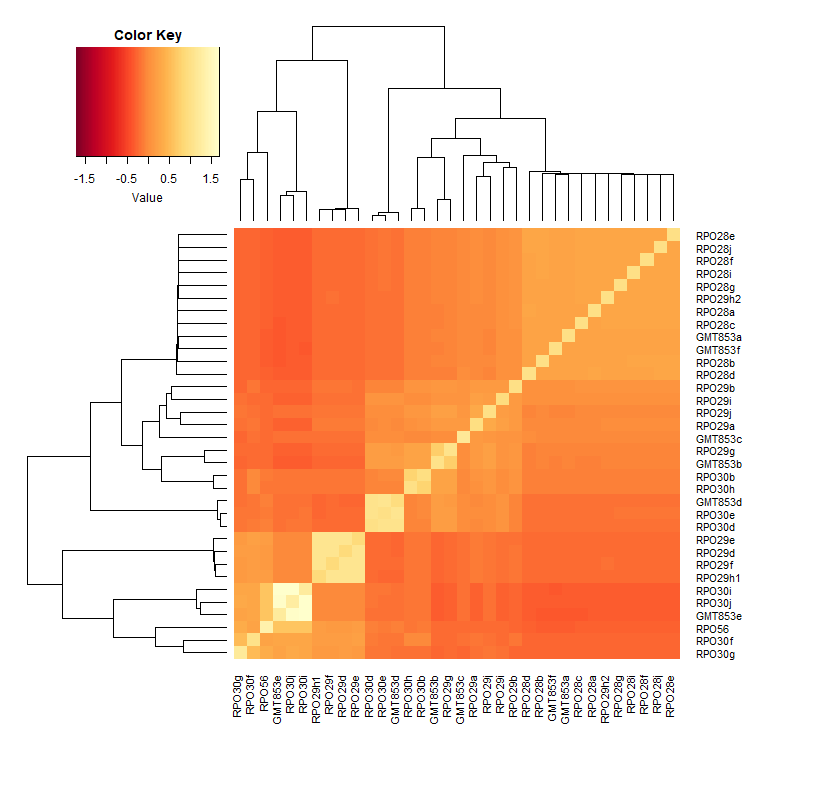

### Online Resource 7 Genomic relationship matrices for all sampled individuals of *P.* sp. Evans Crown. Lighter colours correspond with individuals that are more closely related, with the lightest values at the central diagonal representing the relationship between each specimen and itself. Sample codes follow those outlined in Online Resource 1

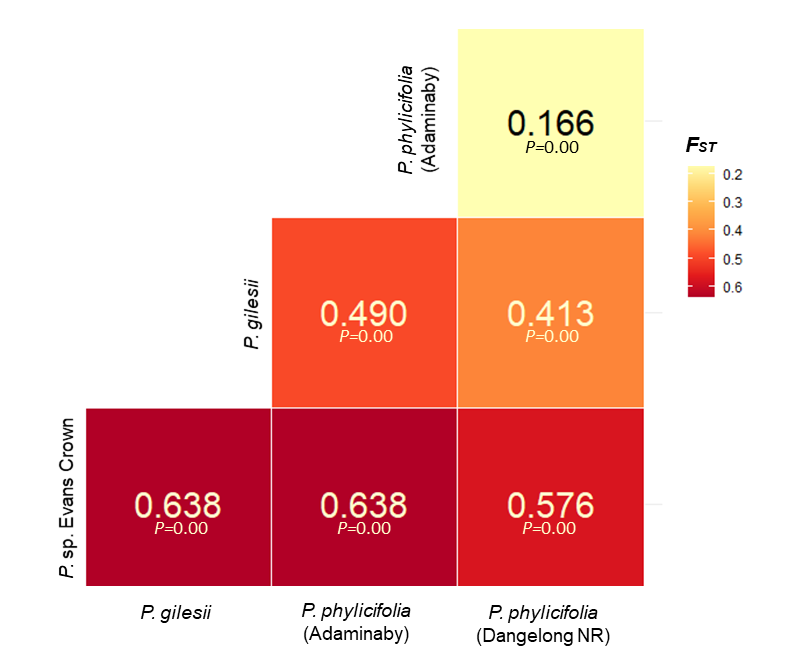

### Online Resource 8 Pairwise *F_ST_* and *P* values between populations with ≥ 5 samples of *P. gilesii*, *P. phylicifolia* and *P.* sp. Evans Crown following removal of putative clones

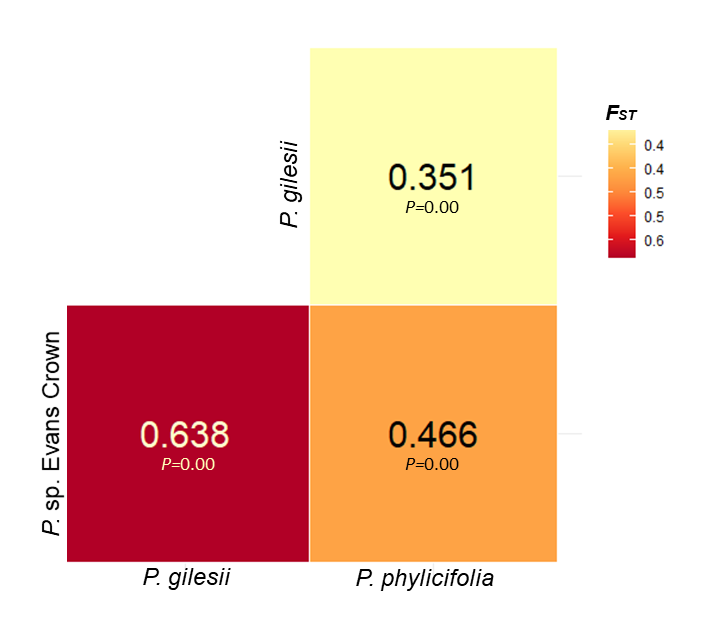

### Online Resource 9 Species-level pairwise *F_ST_* and *P* values between *P. gilesii*, *P. phylicifolia* and *P.* sp. Evans Crown following removal of putative clones

### Online Resource 10 Correlation of characters with ordination pattern (PCC) from semi-strong hybrid multidimensional scaling (SSH MDS) ordination for all selected specimens of *P.* sp. Evans Crown, *P. gilesii* and *P. phylicifolia*. Ad = adaxial surface, Ab = abaxial surface

| **Character #** | **Character** | ***X*** | ***Y*** | ***Z*** | ***r^2^*** |
| --- | --- | --- | --- | --- | --- |
| 10 | Midrib hair direction | -0.386 | -0.612 | 0.69 | 0.971 |
| 19 | Prophyll hair density | -0.551 | -0.112 | 0.827 | 0.966 |
| 22 | Calyx outer surface hair density | -0.666 | -0.164 | 0.727 | 0.929 |
| 11 | Lamina ad hair density | -0.562 | 0.073 | 0.824 | 0.925 |
| 8 | Lamina ab hair density | -0.553 | 0.264 | 0.79 | 0.923 |
| 12 | Ad hair length | -0.691 | -0.35 | 0.633 | 0.922 |
| 14 | a_1_ axis hair density | -0.269 | 0.32 | 0.908 | 0.921 |
| 21 | Calyx outer surface hair direction | -0.742 | -0.279 | 0.61 | 0.918 |
| 4 | Petiole hair direction | 0.068 | -0.11 | 0.992 | 0.912 |
| 26 | Calyx ad lobe length | 0.923 | -0.036 | 0.383 | 0.873 |
| 24 | Calyx ab lobe width | 0.72 | -0.388 | 0.575 | 0.835 |
| 27 | Calyx ad lobe width | 0.759 | -0.208 | 0.617 | 0.827 |
| 23 | Calyx ab lobe length | 0.914 | -0.27 | 0.303 | 0.816 |
| 2 | Branch hair density | -0.14 | -0.463 | 0.875 | 0.805 |
| 9 | Ab hair length | 0.415 | -0.839 | 0.351 | 0.787 |
| 6 | Lamina width | 0.884 | -0.173 | 0.435 | 0.771 |
| 18 | Prophyll L:W ratio | -0.073 | 0.742 | -0.666 | 0.732 |
| 5 | Lamina length | 0.749 | -0.536 | 0.39 | 0.66 |
| 25 | Calyx ab lobe L:W ratio | -0.281 | 0.122 | -0.952 | 0.62 |
| 3 | Branch hair length | 0.768 | 0.156 | 0.621 | 0.614 |
| 28 | Calyx ad lobe L:W ratio | -0.162 | 0.081 | -0.984 | 0.587 |
| 1 | Branch hair direction | -0.506 | 0.812 | 0.291 | 0.58 |
| 16 | Prophyll length | 0.454 | -0.529 | 0.717 | 0.578 |
| 15 | a_1_ axis hair direction | -0.371 | 0.831 | 0.416 | 0.552 |
| 20 | Prophyll hair direction | -0.34 | 0.339 | 0.877 | 0.55 |
| 17 | Prophyll width | -0.111 | -0.842 | 0.528 | 0.431 |
| 29 | Calyx abaxial:adaxial lobe length ratio | -0.069 | -0.887 | -0.457 | 0.43 |
| 13 | a_1_ axis length | 0.945 | 0.062 | -0.322 | 0.416 |
| 7 | Lamina L:W ratio | -0.734 | -0.45 | -0.508 | 0.326 |
